# Blockade of pro-fibrotic response mediated by the miR-143/-145 cluster prevents targeted therapy-induced phenotypic plasticity and resistance in melanoma

**DOI:** 10.1101/2021.07.02.450838

**Authors:** S. Diazzi, A. Baeri, J. Fassy, M. Lecacheur, O. Marin-Bejar, C.A. Girard, L. Lefevre, C. Lacoux, M. Irondelle, C. Mounier, M. Truchi, M. Couralet, A. Carminati, I. Berestjuk, F. Larbret, G. Vassaux, J.-C. Marine, M. Deckert, B. Mari, S. Tartare-Deckert

## Abstract

Lineage dedifferentiation towards a mesenchymal-like state is a common mechanism of adaptive response and resistance to targeted therapy in melanoma. Yet, the transcriptional network driving this phenotypic plasticity remains elusive. Remarkably, this cellular state displays myofibroblast and fibrotic features and escapes MAPK inhibitors (MAPKi) through extracellular matrix (ECM) remodeling activities. Here we show that the anti-fibrotic drug Nintedanib/BIBF1120 is active to normalize the fibrous ECM network, enhance the efficacy of MAPK-targeted therapy and delay tumor relapse in a pre-clinical model of melanoma. We also uncovered the molecular networks that regulate the acquisition of this resistant phenotype and its reversion by Nintedanib, pointing the miR-143/-145 pro-fibrotic cluster as a driver of the therapy-resistant mesenchymal-like phenotype. Upregulation of the miR-143/-145 cluster under BRAFi/MAPKi therapy was observed in melanoma cells *in vitro* and *in vivo* and was associated with an invasive/undifferentiated profile of resistant cells. The 2 mature miRNAs generated from this cluster, miR-143-3p and miR-145-5p collaborated to mediate phenotypic transition towards a drug resistant undifferentiated mesenchymal-like state by targeting Fascin actin-bundling protein 1 (FSCN1), modulating the dynamic crosstalk between the actin cytoskeleton and the ECM through the regulation of focal adhesion dynamics as well as contributing to a fine-tuning of mechanotransduction pathways. Our study brings insights into a novel miRNA-mediated regulatory network that contributes to non-genetic adaptive drug resistance and provides proof-of-principle that preventing MAPKi-induced pro-fibrotic stromal response is a viable therapeutic opportunity for patients on targeted therapy.

## INTRODUCTION

Because of its high mutational burden, metastasis propensity, and resistance to treatment, cutaneous melanoma is one of the most aggressive human cancers and the deadliest form of skin cancer (1). Melanoma is a non-epithelial tumor that originates from neural crest-derived and pigment producing melanocytes in the skin. Genetic alterations in the *BRAF*, *NRAS*, or *NF1* genes define melanoma subtypes and lead to the MAPK pathway hyperactivation (2, 3). Current therapeutic options for BRAF^V600E/K^ metastatic melanoma include MAPK-targeted therapies, which show remarkable efficacy during the first months of treatment (4, 5). However, the majority of patients treated with a combination of BRAF inhibitor (BRAFi) and MEK inhibitor (MEKi) inevitably relapse within months (6). Genetic mechanisms of resistance cannot singly explain the acquisition of therapy resistance in melanoma and non-genetic heterogeneity actively participates in drug tolerance (7, 8). Extensive studies have been carried out to dissect the non-mutational mechanisms of resistance (9, 10). Genetic and non-genetic mechanisms of resistance are frequently linked and not mutually exclusive (8). Non-genetic resistance is due to the intrinsic melanoma cell phenotypic plasticity, i.e., ability to undergo transcriptional and epigenetic reprogramming in response to environmental challenges or upon therapy (11). These adaptive mechanisms exploit the developmental plasticity of melanoma cells and often result in an undifferentiated state characterized by upregulation of receptor tyrosine kinases (RTK) like AXL, downregulation of melanocyte differentiation transcription factors MITF and SOX10 (12) and acquisition of mesenchymal and invasive features (9, 10, 13–18).

Tumors are shaped dynamically by reciprocal crosstalk between cancer cells and the ECM through cellular-ECM interactions and stromal matrix remodeling. Recent findings indicated that elevated ECM production and remodeling contribute to adaptive and acquired resistance to BRAFi therapy by conferring a drug-protective niche to melanoma cells (19–22). Moreover, we recently reported that undifferentiated mesenchymal-like BRAFi-resistant cells exhibit myofibroblast/cancer associated fibroblast (CAF)-like features leading to pro-fibrotic ECM reprogramming *in vitro* and *in vivo* (22, 23). Cell autonomous ECM deposition and remodeling abilities adopted by melanoma cells after MAPKi treatment results in cross-linked collagen matrix and tumor stiffening fostering a feedforward loop dependent on the mechanotransducers YAP and MRTFA and leading to therapy resistance (22). Thus, this pro-fibrotic-like response, typical of the early adaptation and acquired resistance to MAPK inhibition, provides a therapeutic escape route through the activation of alternative survival pathways mediated by cell-matrix communications. However, the signaling networks underlying the acquisition of this undifferentiated, mesenchymal-like melanoma cell state and drug resistant behavior remain unclear.

We reasoned that therapeutic approaches aimed at preventing this targeted therapy-induced abnormal pro-fibrotic-like response could represent rationale combination strategies to normalize the fibrous stroma and overcome non-genetic resistance in BRAF^V600E^-mutant melanomas. We show here that the anti-fibrotic drug Nintedanib (BIBF1120) improves the response of the BRAFi/MEKi targeted therapy in a pre-clinical model of melanoma as well as in BRAF-mutated cell lines by preventing MAPKi-induced lineage dedifferentiation, ECM reprogramming and mesenchymal traits. We also identified the master regulator associated with the acquisition of this pro-fibrotic and dedifferentiation program, pointing the miR-143/-145 cluster as a driver of the phenotype switching to a drug resistant mesenchymal-like cell state.

## RESULTS

### Nintedanib/BIBF1120 prevents MAPKi-induced pro-fibrotic-like response, enhances targeted therapy efficiency and delays tumor relapse

In order to limit ECM reprogramming and collagen remodeling associated with therapy resistance and relapse in melanoma, we tested the effect of the anti-fibrotic drug Nintedanib (BIBF1120), a triple inhibitor of PDGFR, VEGFR and FGFR used to treat idiopathic pulmonary fibrosis (IPF) in combination with BRAFi/MEKi in a syngeneic model of transplanted murine YUMM1.7 Braf-mutant melanoma (24). YUMM1.7 cells were subcutaneously injected and tumors were treated with vehicle, BIBF1120 alone, a combination of BRAFi plus MEKi, or the triple combination (Fig. 1A). BIBF1120 did not display any anti-melanoma effect when administered alone, slightly slowing down tumor growth but not triggering tumor volume decrease. Administration of the BRAFi/MEKi initially reduced tumor growth but after three weeks of treatment, tumor growth resumed and 100% of tumors relapsed. Importantly, combination of MAPK-targeted therapies and BIBF1120 significantly delayed relapse and led to complete remission in 33% of mice (2 out of 6) (Fig. 1B-C and Supplementary Fig. 1A). Overall, the combined treatment significantly improved mice survival (Fig. 1C) without body weight loss or sign of toxicity throughout the study (Fig. 1D). As previously described in melanoma xenograft models (22), an extensive deposition of collagens and increased expression of ECM remodeling and myofibroblast markers were observed in YUMM1.7 tumors treated with the combination BRAFi/MEKi as revealed by picrosirius red staining of collagen fibers and qPCR analysis of typical molecular markers of tumor fibrosis. This response was significantly reduced by the co-administration of BIBF1120 (Fig. 1E-G and Supplementary Fig. 1B). Thus, combination of targeted therapy with the anti-fibrotic drug Nintedanib prevents the appearance of a pro-fibrotic matrix observed upon MAPK-targeted therapy exposure and significantly delays the onset of resistance *in vivo*.

**Fig. 1.**
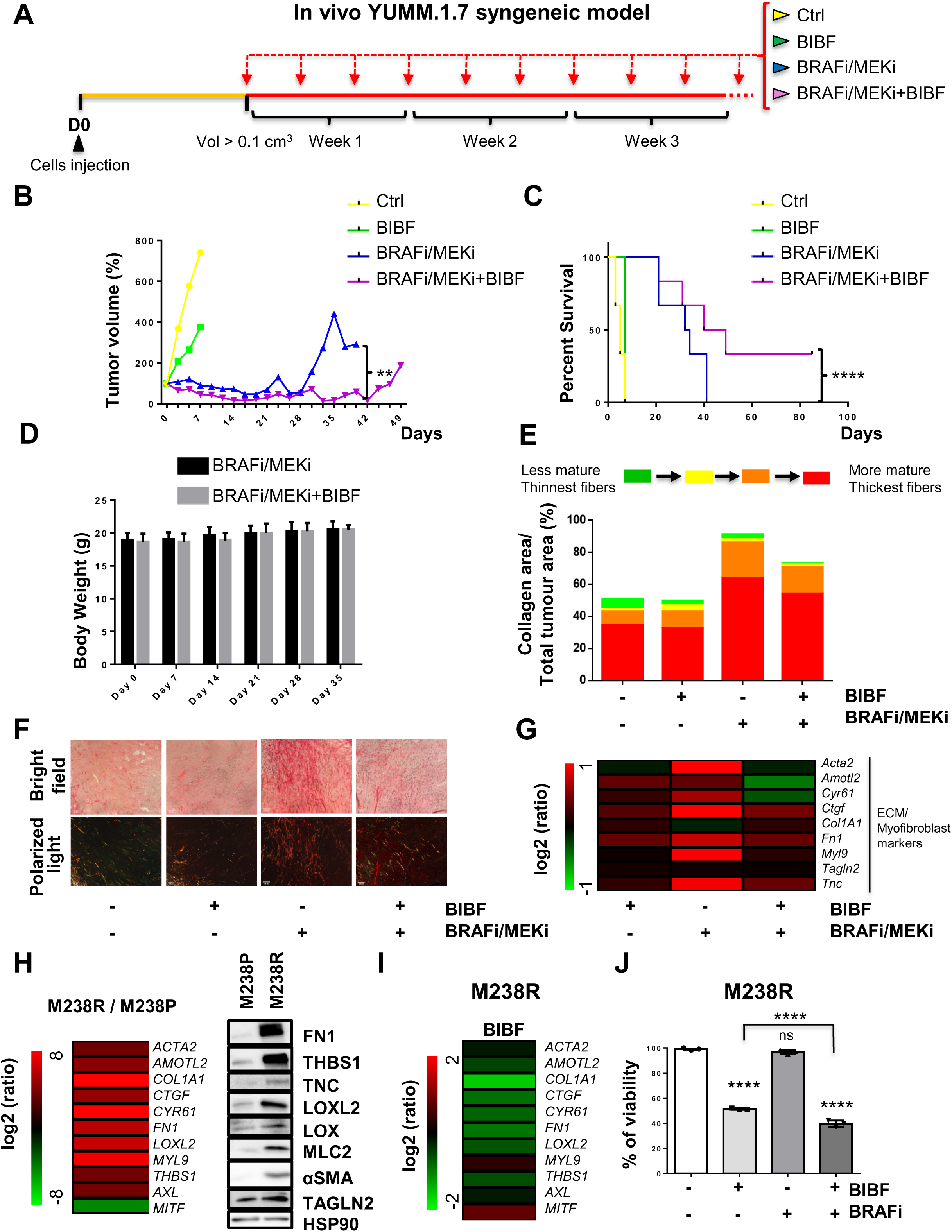
Nintedanib/BIBF1120 prevents MAPKi-induced ECM remodeling, decreases resistance to targeted therapy and delays tumor relapse. (A-G) Mouse YUMM1.7 melanoma cells were subcutaneously inoculated into C57BL/6 mice and when tumors reached 100 mm^3^ mice were treated with vehicle (Ctrl), BIBF1120 (BIBF), MAPK-targeted therapies (BRAFi, Vemurafenib + MEKi, Trametinib) or MAPK-targeted therapies plus BIBF. **(B)** Representative median graphics showing tumor growth following treatment. Two-way ANOVA was used for statistical analysis. ** P≤0.01. **(C)** Kaplan-Meier survival curves of mice treated with the indicated therapies. Log rank (Mantel-Cox) statistical test was used for MAPK-targeted therapies vs MAPK-targeted therapies/BIBF1120. ****P≤0.0001. **(D)** Mouse body weight was measured at the indicated times. Data shown are mean±SD (n=6). **(E-F)** Tumor sections were stained with picrosirius red and imaged under polarized light. **(E)** Quantification of collagen fibers thickness. **(F)** Representative image of collagen fibers network in tumors from mice under the different treatments. Scale bar 100 μM. **(G)** Heatmap showing the differential expression of ECM and myofibroblast/CAF genes in mice treated with MAPK-targeted therapies with or without BIBF compared to control mice (log2 ratio, n=5). **(H)** Heatmap and Western Blot showing the expression of ECM, myofibroblast/CAF and phenotype switch markers in human M238R cells compared to the parental M238P cells. Heatmap represents the mean of expression of 3 independent experiments by RT-qPCR. **(I)** Heatmap showing the expression of ECM, myofibroblast/CAF and phenotype switch markers in human M238R cells treated with BIBF1120 (BIBF) (2 µM, 72 h) or vehicle alone by RT-qPCR (n=3). **(J)** Crystal violet viability assay of M238R cells treated with BRAFi (Vemurafenib, 3 μM), BIBF1120 (BIBF, 2 μM) or with BRAFi (3 μM) plus BIBF (2 μM) for 72 h. Paired Student t-test was used for statistical analysis. ****P≤0.0001. Data is represented as mean ± SD from a triplicate representative of 3 independent experiments.

We next examined the impact of Nintedanib on ECM reprogramming and cell phenotype switching in the context of early adaptation and resistance to MAPK targeted therapy in human BRAF^V600E^ mutated melanoma M238P cells. BIBF1120 strongly attenuated targeted drugs-induced ECM/myofibroblast-related signatures, prevented the undifferentiated AXL^high^ MITF^low^ phenotype switch (Supplementary Fig. 1C) and potentiated the effect of the BRAFi/MEKi cocktail on M238P cell viability (Supplementary Fig. 1D). The efficacy of the described treatment to reduce upregulation of Fibronectin (FN1) and LOXL2 expression was confirmed at protein levels by Western Blot analysis. Of note, a strong activation of AKT induced by the BRAFi/MEKi cocktail was fully inhibited by BIBF1120, suggesting that the anti-fibrotic drug is able to counteract the rewiring of alternative survival pathway observed upon MAPK oncogenic pathway inhibition (Supplementary Fig. 1E) (17).

We finally evaluated the effect of BIBF1120 on the undifferentiated mesenchymal-like resistant M238R cells obtained through chronic exposure of the M238P cells to the BRAFi Vemurafenib (17). We recently demonstrated that this resistant cell line exhibits low expression of the differentiation factor MITF and high AXL levels and displays a strong myofibroblast-like phenotype with expression of classical ECM and contractile markers such as smooth muscle actin-α (αSMA) and Myosin light chain 2 (MLC2) as well as ECM remodeling activities compared with parental M238P cells (Fig. 1H) (22). BIBF1120 was able to attenuate melanoma undifferentiated state markers and expression of ECM and myofibroblast/CAF-related signature (Fig. 1I), but also significantly decreased cell viability and resistance to BRAFi (Fig. 1J). These findings indicate that an anti-fibrotic therapy is able to revert the undifferentiated-mesenchymal resistant phenotype and potentiate targeted therapy in human melanoma cells.

### Suppression of MAPKi-induced resistant pro-fibrotic phenotype by Nintedanib is associated with loss of miR-143/145 cluster expression

Next we investigated the molecular mechanisms associated with the emergence of MAPKi-induced mesenchymal and pro-fibrotic phenotype and its inhibition by Nintedanib/BIBF1120. Because several microRNAs (miRNAs), named FibromiRs, have been shown to play key roles in the initiation and progression of fibrotic processes in various organs (25–28), we performed an expression screening to compare the level of these FibromiRs in BRAF^V600E^ mutant melanoma cells sensitive to MAPK-targeted therapies (M229P, M238P, M249P) compared to their corresponding resistant counterparts (17). The screening identified miR-143-3p and miR-145-5p, localized within the miR-143/145 cluster on chromosome 5 as the best hits with a strong upregulation in AXL^high^ MITF^low^ mesenchymal-like resistant M238R and M229R cells tested compared to parental cells (Fig. 2A, Supplementary Fig. 2A). Similar results were obtained in the mesenchymal resistant UACC62R cells (29) (Supplementary Fig. 2A). In contrast, acquisition of resistance through secondary NRAS mutation was not associated with increased expression of miR-143-3p and miR-145-5p in the non-mesenchymal AXL^low^ MITF^high^ M249R cells (Fig. 2A, Supplementary Fig. 2A). We next examined whether a treatment with BRAFi, MEKi, or a combination of both was able to modulate the expression of the cluster. The two drugs, alone or in combination, significantly increased miR-143-3p and miR-145-5p expression levels in all BRAF^V600E^ mutant melanoma cells tested including patient-derived short-term melanoma cultures (Supplementary Fig. 2B-F). This strong induction was abolished when the BRAFi/MEKi treatment was combined with BIBF1120, both in melanoma cell lines cultured *in vitro* (Fig. 2B) and in the YUMM.1.7 syngeneic model (Fig. 2C) presented in Fig.1. Overall, the expression of the miR-143/-145 cluster paralleled the phenotypic switch associated with a mesenchymal resistant phenotype.

**Fig. 2.**
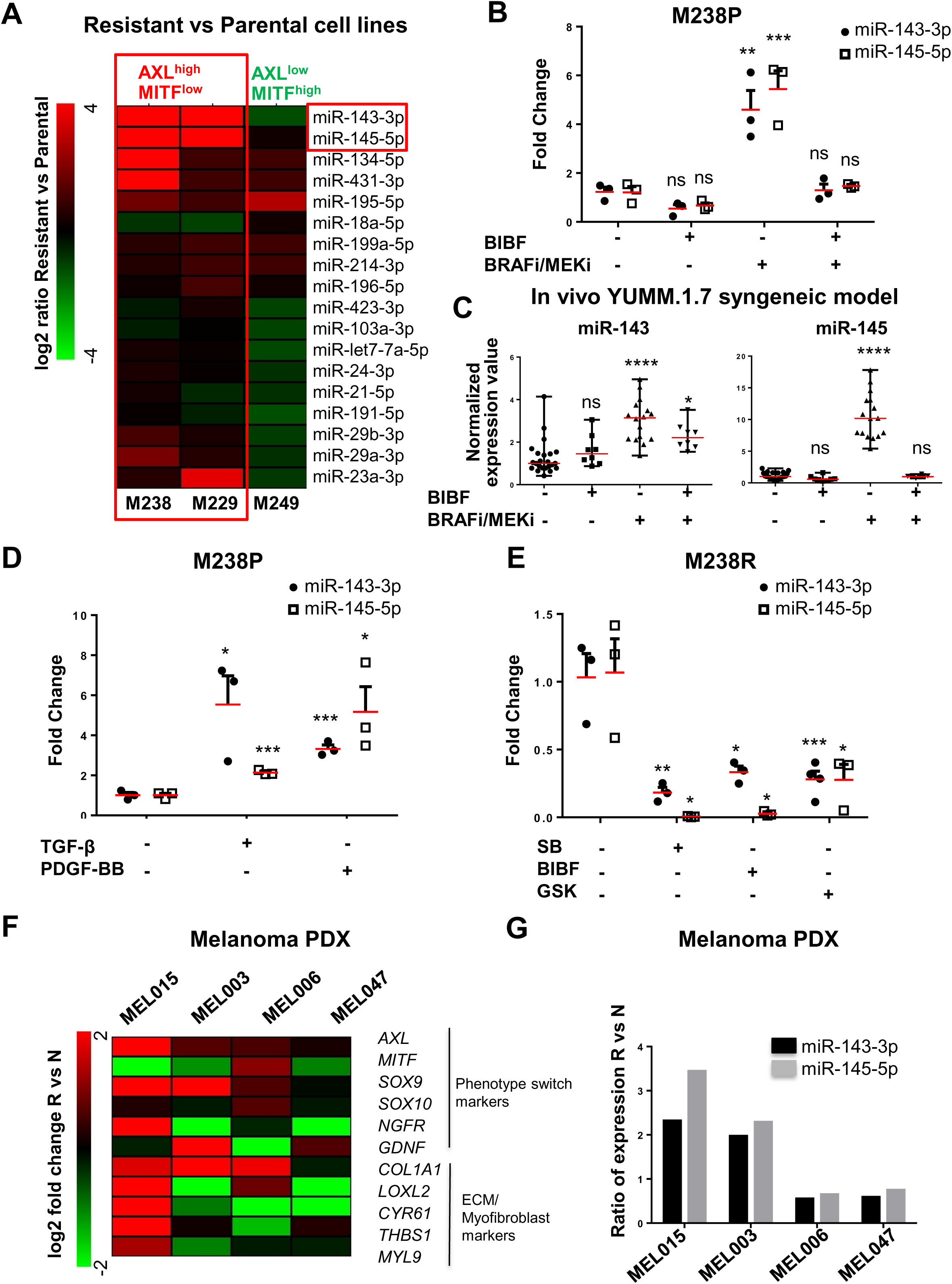
Expression of miR-143/-145 is correlated with the undifferentiated mesenchymal-like MAPKi-resistant phenotype and is negatively regulated by Nintedanib/BIBF1120. **(A)** Heatmap showing the differential expression of a selection of miRNAs known as “FibromiRs” in human BRAF^V600E^ mutant melanoma cells sensitive to MAPK-targeted therapies (M229, M238, M249) and the corresponding BRAFi-resistant cells. The type of resistance for each cell line is indicated. Expression was evaluated by RT-qPCR; log2 (R/P) is indicated for each cell line. **(B)** Relative miRNA expression levels was quantified in M238P cells treated for 72 h with BIBF1120 (BIBF, 2 μM), MAPK-targeted therapies (BRAFi, Vemurafenib + MEKi, Trametinib) (1µM), or with MAPK-targeted therapies (1 μM) plus BIBF (2µM) by RT-qPCR and normalized to miR-16-5p. Data is represented as mean ± SEM from a triplicate representative of 3 independent experiments. One-way Anova was used for statistical analysis. **P≤0.01, ***P≤0.001. **(C)** Expression of miR-143-3p and miR-145-5p in control mice and mice treated with the indicated therapies (see legend of Fig.1 for details) was quantified by RT-qPCR. One-way Anova was used for statistical analysis. *P≤0.05, ****P≤0.0001. **(D)** Relative miRNA expression levels was quantified in M238P cells stimulated for 48h with TGF-β (10 ng/mL) or PDGF-BB (20 ng/mL) by RT-qPCR and normalized to miR-16-5p. Data is represented as mean ± SEM from a triplicate representative of 3 independent experiments. P-values were calculated using Paired Student t-test. *P≤0.05, ***P≤0.001. **(E)** Relative miRNA expression levels was quantified in M238R cells treated for 48 h with the triple kinase inhibitor Nintedanib/BIBF1120 (BIBF, 2 μM), the TGF-β receptor kinase inhibitor SB431542 (SB, 10 µM), and the pan-AKT inhibitor GSK690693 (GSK, 10 µM) by RT-qPCR. Data is represented as mean ± SEM from a triplicate representative of 3 independent experiments. P-values were calculated using Paired Student t-test. *P≤0.05, **P≤0.01; ***P≤0.001. **(F, G)** Phenotype switch/invasive/ECM markers **(F)** and relative miRNAs expression levels **(G)** were quantified in therapy-naïve (N) and resistant (R) PDX samples. The log2 fold change or the ratio of the fold change R vs N is shown for each couple of samples.

Given the critical role of RTKs upregulation such as PDGFR and of the pro-fibrotic TGF-β signaling pathway overactivation in mesenchymal resistance (12, 17, 22, 23), we stimulated MAPKi sensitive melanoma cells with PDGF-BB or with TGF-β and analyzed miR-143-3p and miR-145-5p expression. Both TGF-β and PDGF-BB triggered a strong upregulation of miR-143/-145 expression in M238P cells (Fig. 2D). Conversely, treatment of mesenchymal BRAFi-resistant M238R cells with BIBF1120 but also with the TGF-β receptor inhibitor SB431542 and the pan-AKT inhibitor GSK690693 significantly decreased the expression of the two mature miRNAs (Fig. 2E), indicating that both PDGF and TGF-β pathways control the expression of the miR-143/-145 cluster in melanoma cells.

Finally, we investigated the expression of these miRNAs in several Patient-Derived Xenograft (PDX) samples that acquired resistance to BRAFi/MEKi combo-therapy and exhibited distinct phenotypic and molecular profiles. (Fig. 2F) (30). Upregulation of miR-143/-145 cluster between therapy naïve and resistant cells was observed in two different PDX samples, MEL015 and MEL003, with a predominant invasive/undifferentiated transcriptome profile (Fig. 2F-G) (30). The MEL015 resistant model also presented elevated expression of ECM remodeling, myofibroblast and pro-fibrotic markers such as *COL1A1*, *LOXL2*, *CYR61*, *THBS1* and *MYL9*. In contrast, we did not observe an upregulation of the cluster in drug resistant lesions from the two additional PDX models, MEL006 and MEL047, in which the mesenchymal-like signature is not overrepresented (Fig. 2F-G). These data indicate that upregulation of the pro-fibrotic miR-143/-145 cluster is also observed in PDX MAPKi resistant melanomas associated with an invasive/undifferentiated transcriptome profile.

### miR-143/-145 cluster promotes melanoma cell dedifferentiation towards a pro-fibrotic mesenchymal-like state and resistance to MAPK therapeutics

To confirm a potential link between the miR-143/-145 cluster and ECM reprogramming, we first used a gain of function approach consisting in the transient overexpression of miR-143-3p or miR-145-5p in various therapy-naïve BRAF-mutant melanoma cells (Supplementary Fig. 3A). The results showed increased expression of transcripts related to ECM structure and remodeling as well as myofibroblast/CAF markers in cells overexpressing either miRNA compared to miR-Neg control cells (Fig. 3A). Conversely, we next tested whether miR-143-3p or miR-145-5p inhibition can reverse the phenotypic pro-fibrotic response induced by oncogenic BRAF inhibition in M238P melanoma cells. BRAFi treatment was combined with Locked nucleic acid (LNA)-modified antisense oligonucleotides (ASOs) designed against miR-143 (LNA-143), miR-145 (LNA-145) or a control LNA ASO (LNA-Ctrl). RT-qPCR analysis showed that the BRAFi-induced ECM- and myofibroblast/CAF-related gene signature was significant inhibited by LNA-143 and LNA-145 ASOs (Fig. 3B). These results were confirmed at protein level by Western Blot analysis of cell lysates and conditioned media of ECM proteins and cross-linking enzymes as well as myofibroblast/CAF markers using same gain- or loss-of-function approaches (Fig. 3C-D and Supplementary Fig. 3B).

**Fig. 3.**
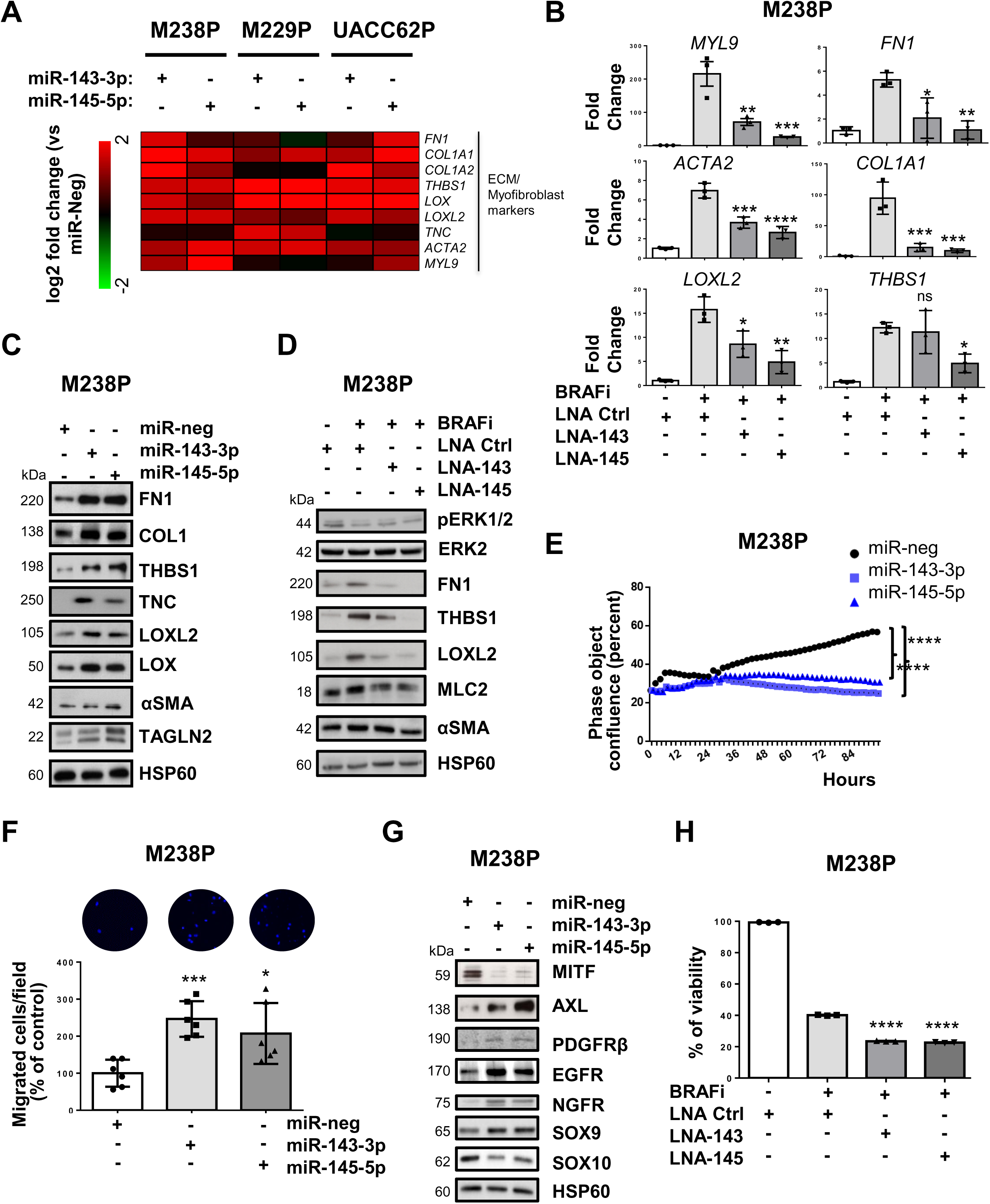
miR-143/-145 cluster promotes ECM reprogramming, melanoma cell dedifferentiation and drug resistance. **(A)** Heatmap showing the differential expression of a selection of ECM-related genes, cytoskeleton and myofibroblast markers in 3 distinct cell lines (M238P, UACC62P, M229P) transfected with the indicated mimics (control (miR-neg), miR-143 or miR-145 mimics, 72 h, 30 nM), assessed by RT-qPCR. (n=3) **(B)** M238P cells were treated 72 h with BRAFi (Vemurafenib,1 μM, 72 h) in the presence or the absence of LNA-based anti-miR-143 (LNA-143) or anti-miR-145 (LNA-145) (50 nM, 72 h). ECM markers RT-qPCR data is represented as mean ± SD from a triplicate representative of at least 3 independent experiments. One-way Anova was used for statistical analysis. *P≤0.05, **P≤0.01, ***P≤0.001, ****P≤0.0001. **(C)** ImmunoBlot analysis of ECM remodeling markers on total cell lysates from parental cells (M238P) transfected as in **(A)**. **(D)** ImmunoBlot analysis of ECM remodeling markers of cells treated with the indicated combination of inhibitors. **(E)** Proliferation curves using time-lapse analysis of cells with the IncuCyte system. Graph shows quantification of cell confluence. 2-way ANOVA analysis was used for statistical analysis. ****P≤0.0001. **(F)** Migration assay performed in Boyden chambers. Representative images showing migrating cells in the different conditions. The histogram represents the quantitative determination of data obtained using ImageJ software. Paired Student t-test was used for statistical analysis. * P≤0.05 ***P≤0.001. **(G)** ImmunoBlot analysis of phenotype switch markers on lysates from cells treated as in **(A)**. **(H)** Crystal violet viability assay of M238P cells treated with the different combinations of inhibitors. Data is represented as mean ± SD from a triplicate representative of at least 3 independent experiments. One-way ANOVA was used for statistical analysis. ****P≤0.0001.

We next investigated whether the cluster contributed to the acquisition of the slow cycling, undifferentiated and invasive cell state. Melanoma cells experienced reduced cell proliferation after ectopic expression of miR-143-3p or miR-145-5p as visualized by Western Blot analysis of cell cycle markers (Supplementary Fig. 4A) and by analysis of cell confluence by live-cell imaging (Fig. 3E and Supplementary Fig. 4B), with an accumulation of cells in the G2/M phase (Supplementary Fig. 4C). Inhibition of proliferation was also accompanied by enhancement of cell migratory abilities, as shown using Boyden chamber assays (Fig. 3F and Supplementary Fig. 4D) as well as by the acquisition of an undifferentiated phenotype, with decreased levels of MITF and SOX10, and increased levels of AXL, PDGFR, EGFR, NGFR and SOX9 (Fig. 3G and Supplementary Fig. 4E). Lentivirus-mediated stable overexpression of the two miRNAs in two distinct melanoma cell lines reproduced increased ECM protein production, inhibition of cell proliferation and transition to an undifferentiated/invasive phenotype (Supplementary Fig. 5A-E) observed upon transient transfection. Acquisition of this features was also linked to a decreased intrinsic sensitivity to MAPKi treatment, as measured by crystal violet survival assays performed on melanoma cells stably overexpressing miR-143/-145 cluster compared to control cells (Supplementary Fig. 5F). Conversely, targeting the two miRNAs by ASOs in combination with BRAFi improved the efficacy of the targeted therapy (Fig. 3H, Supplementary Fig. 6A-B), demonstrating that miR-143/-145 cluster upregulation in response to BRAF^V600E^ pathway inhibition represents a pivotal adaptive resistance mechanism to MAPK therapeutics.

### Identification of miR-143-3p / miR-145-5p targets functionally associated with the undifferentiated mesenchymal-like phenotype in melanoma cells

To identify miR-143-3p and miR-145-5p targets associated with the resistant mesenchymal phenotype, we first combined *in silico* target prediction tools and experimental transcriptomic approaches using the miRonTop web tool (31) in M238R or M238P cells following transient transfection of mimics (Fig. 4A-B) or stable lentivirus transduction. Functional annotation of the gene expression profiles associated with miRNA overexpression showed a strong overlap in pathways associated with cell migration and invasion, cell cycle as well as cytoskeleton organization (Supplementary Table 1). The predicted targets for each of the mature miRNAs were significantly overrepresented among the downregulated genes in response to the corresponding mimics transfection (Fig. 4B). A first set of target candidates were identified by crossing these predicted targets and the genes shown experimentally to be downregulated in resistant M238R cells compared to parental M238P cells (Fig. 4C).

**Fig. 4.**
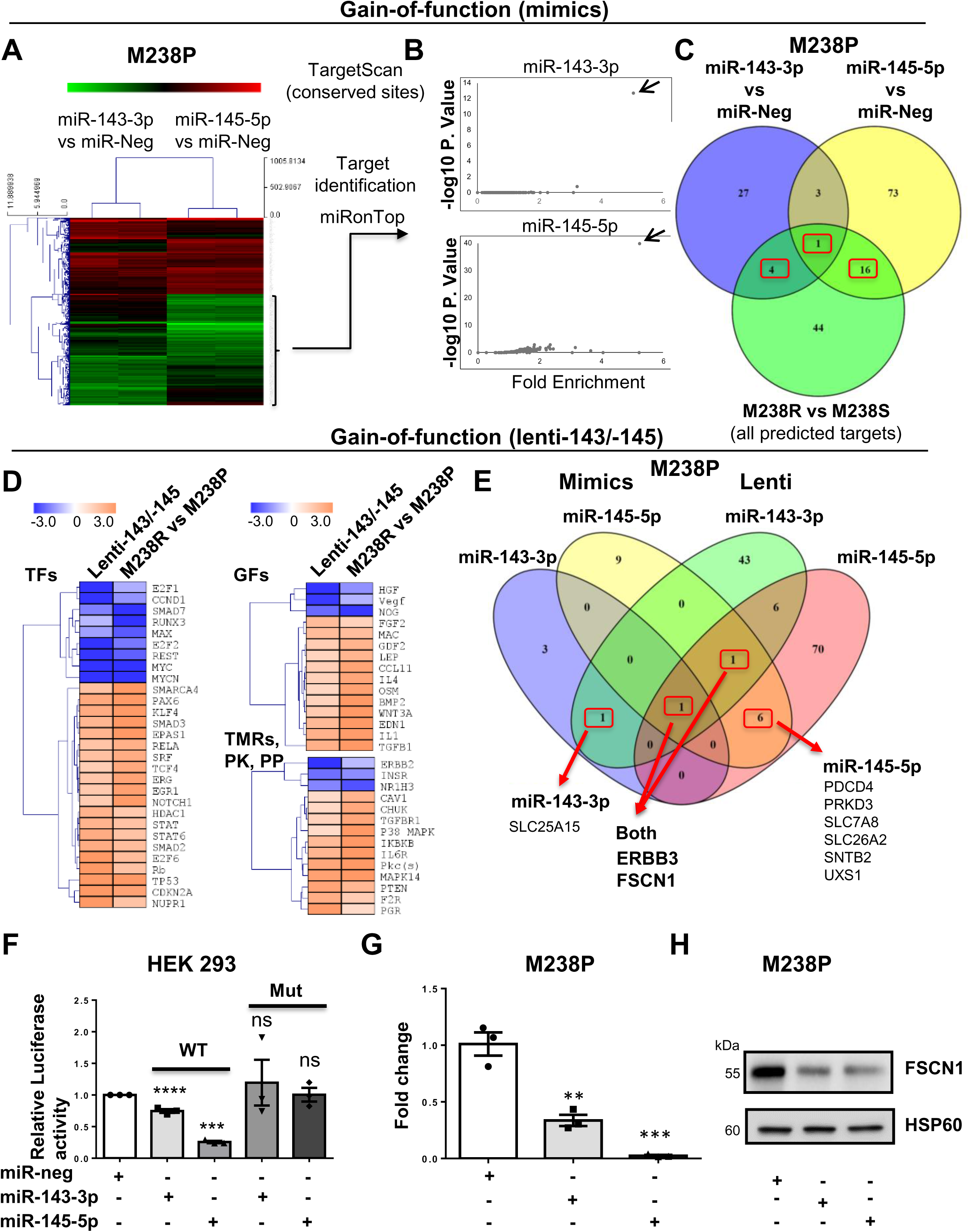
Identification of gene targets and cellular pathways functionally associated with the miR-143/-145 cluster-mediated undifferentiated/mesenchymal-like melanoma cell phenotype. (A-C) M238P cells were transfected separately with miR-143-3p, miR-145-5p or a negative control (miR-neg) mimics and RNA content was analyzed using whole genome microarrays (dataset 1, n=2). **(A)** Heatmap showing the genes differentially expressed after individual miRNA mimic overexpression. **(B)** Overrepresentation of miRNA predicted targets in the set of downregulated transcripts following miR-143-3p and miR-145-5p mimics transfection using miRonTop webtool. Each arrow indicates the corresponding overexpressed miRNA. **(C)** Venn diagram showing the selection of the best target candidates (red boxes) using miR-143-3p and miR-145-5p mimics transfection as well as comparison of M238R and M238P transcriptomic profiles. **(D-E)** M238P cells were transduced with a miR-143/-145 construct and selected for stable expression of the cluster or transduced with a control vector, followed by RNA-seq analysis (dataset 2, n=2). **(D)** Heatmap highlighting the common predicted upstream regulators altered in cells stably expressing the miR-143/-145 cluster and M238R cells compared to control M238P cells. A subset of common regulators (out of the top 50 scores) corresponding to transcription factors (TFs), cytokines and growth factors (GFs), transmembrane receptors, kinases and phosphatases is shown. Red arrows indicate annotations related to the TGF-β pathway. **(E)** Venn diagram summarizing the comparison of the best-predicted targets following the 2 gain-of-function approaches. Subsets of miR-143-3p and miR-145-5p predicted targets downregulated by both mimics and stable lentivirus expression are shown (red boxes). **(F)** Luciferase assay in HEK cells overexpressing miR-143 or miR-145 transfected with a plasmid harboring the WT or muted sequence of the miR-143 and miR-145 binding sites present in FSCN1 3’UTR. Each bar represents the mean ± SE of experiments performed at least in triplicate. ***P≤0.001 ****P≤0.0001. P-values were calculated using Paired Student t-test. **(G)** RT-qPCR analysis of FSCN1 expression in parental cells (M238P) transfected with the indicated mimics. Data is represented as mean ± SE from a triplicate representative of at least 3 independent experiments. Paired Student t-test was used for statistical analysis. ** P≤0.01 ***P≤0.001. **(H)** Western Blot analysis of FSCN1 expression in parental cells (M238P) transfected with the indicated mimics.

Second, RNAs from cells stably overexpressing the miR-143/-145 cluster were analyzed by RNA-sequencing and processed through Ingenuity Pathway Analysis (IPA) to identify the common regulators (transcription factors, growth factors, cytokines, transmembrane receptors, kinases, and phosphatases) between parental cells overexpressing the cluster and resistant cells (Fig. 4D). These analyses notably highlighted changes related to decreased cell proliferation, increased cell invasion and fibrotic pathways activation. To narrow the best target candidates, we finally compared the best-predicted targets based on the two different gain-of-function approaches (Supplementary Table 2 and 3). This strategy resulted in selecting one target candidate for miR-143-3p, 6 target candidates for miR-145-5p and 2 target candidates for both miR-143-3p and miR-145-5p (Fig. 4E). We started with investigations on the F-acting bundling protein Fascin1 (FSCN1), a key regulator of cytoskeleton dynamics, previously associated with tumor growth, migration, invasion and metastasis (32). Using long-reads Nanopore sequencing data, we confirmed lower levels of FSCN1 transcript in M238R compared with M238P cells while reads corresponding to the putative miR-143/-145 cluster primary transcript could be only detected in M238R cells (Supplementary Fig. 7A). The characterization of hFSCN1 3’UTR sequence revealed the presence of 2 miR-143-3p and 4 miR-145-5p binding sites. Validation of these sites was first performed using a luciferase reporter corresponding to the full 3’UTR FSCN1 harboring WT or a mutated sequence of the miRNA recognition elements (Fig. 4F and Supplementary Fig. 7B). Finally, qPCR and Western Blot analyses confirmed that FSCN1 was downregulated at both mRNA and protein levels upon miR-143-3p and miR-145-5p ectopic expression in various melanoma cells as well as in cells stably overexpressing the cluster (Fig. 4G-H and Supplementary Fig. 7C-D).

### FSCN1 is a functional miR-143/-145 target contributing to the phenotypic switch towards the undifferentiated / mesenchymal-like and resistant state

Considering the strong expression of the miR-143/-145 in BRAF^V600E^ mutant mesenchymal-like resistant cells compared to their parental counterparts, we compared FSCN1 expression levels in various pairs of resistant and sensitive melanoma cell lines. Western blot indicated that FSCN1 protein levels were lower in undifferentiated mesenchymal resistant cells compared to parental cells, while on the other hand they were elevated in M249R melanoma cells acquiring genetic resistance compared to parental cells (Supplementary Fig. 8A). We then confirmed the opposite regulation of FSCN1 and miR-143/-145 cluster expression upon BRAFi treatment both *in vivo* using xenografted nude mice and *in vitro* with different human BRAF mutant melanoma cells (Fig. 5A and Supplementary Fig. 8B). Finally, FSCN1 levels were partially restored in M238P cells treated with BRAFi when Vemurafenib was combined with the LNA-miR-143 or LNA-miR-145, as visualized by immunofluorescence staining (Fig. 5B), suggesting that FSCN1 downregulation upon BRAFi exposure is due to increased expression of miR-143-3p and miR-145-5p.

**Fig. 5.**
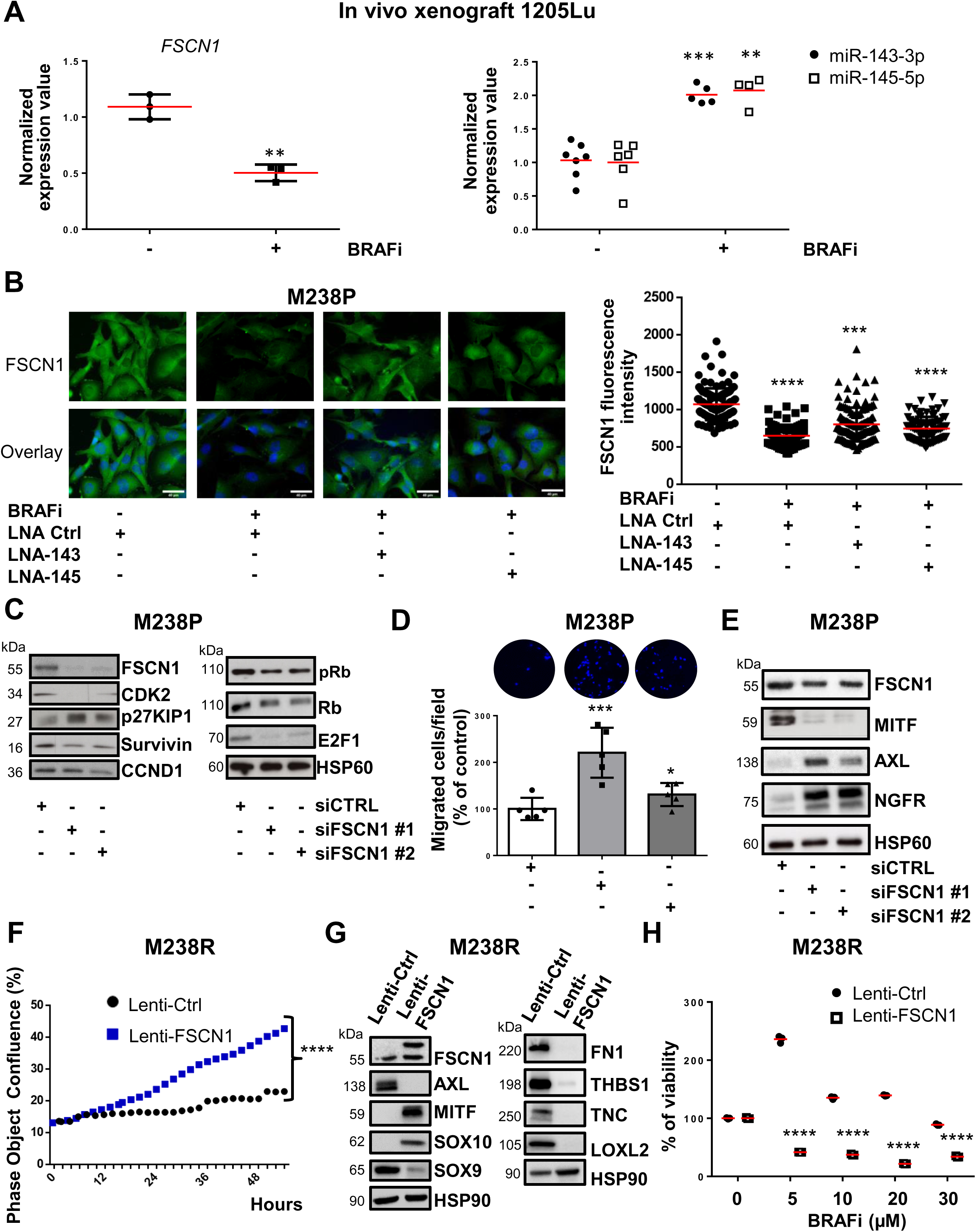
FSCN1 is a functional miR-143/-145 target contributing to the phenotypic switch towards the undifferentiated/mesenchymal-like state. **(A)** qPCR analysis of FSCN1, miR-143 and miR-145 expression in a 1205Lu xenograft nude mice model treated with the BRAFi Vemurafenib compared to control mice. Paired Student t-test has been used for statistical analysis. **P≤0.01, ***P≤0.001. **(B)** FSCN1 immunofluorescent staining and quantification of fluorescence intensity in cells (M238P) treated with the different combinations of inhibitors. Mann-Whitney U test has been used for statistical analysis. ***P≤0.001, ****P≤0.0001. Scale bar 40 μM. **(C-E)** M238P cells were transfected with two different sequences of siRNAs vs FSCN1 or with a control siRNA (72 h, 100 nM). **(C)** ImmunoBlot analysis of cell cycle markers on cell lysates from M238P cells cultured for 72 h following transfection with the different siRNAs. **(D)** Migration assay performed in Boyden chambers. Representative images showing migration of M238P cells treated with the indicated siRNAs. The histogram represents the quantitative determination of data obtained using ImageJ software. Paired Student t-test was used for statistical analysis. *P≤0.05, ***P≤0.001 **(E)** ImmunoBlot analysis of phenotype-switch markers on cell lysates from M238P cells transfected with the indicated siRNAs. **(F-H)** BRAFi-resistant M238R cells overexpressing FSCN1 were obtained after transduction with a FSCN1 lentiviral construct. M238R transduced with a Ctrl lentivirus were used as control. **(F)** Effect of FSCN1 overexpression on cell proliferation assessed by time-lapse analysis using the IncuCyte system. Graph shows quantification of cell confluence. 2-way ANOVA analysis was used for statistical analysis. ****P≤0.0001. **(G)** ImmunoBlot analysis of FSCN1, phenotype-switch markers and ECM remodeling markers on cell lysates from control and FSCN1 overexpressing cells. **(H)** Crystal violet viability assay of M238R cells stably overexpressing FSCN1 treated with the BRAFi Vemurafenib. (6 days, Vemurafenib (Vemu) 5, 10, 20 or 30 μM). Paired Student t-test was used for statistical analysis. Data is represented as mean ± SD from a triplicate representative of at least 3 independent experiments. ****P≤0.0001.

To evaluate the influence of FSCN1 downregulation among the various cellular effects mediated by miR-143-3p and miR-145-5p, we then performed a loss-of-function experiment using FSCN1 specific siRNAs in BRAF-mutant parental melanoma cells. Western Blot analysis of cell cycle markers (Fig. 5C and Supplementary Fig. 8C) and cell confluence analysis by live-cell imaging (Supplementary Fig. 8D) showed reduced proliferation after downregulation of FSCN1. This slow-cycling state induced by FSCN1 silencing was accompanied by an enhancement in cell migratory abilities (Fig. 5D and Supplementary Fig. 8E). Moreover, FSCN1 invalidation modulated melanoma cells differentiation state, inducing the switch to a poorly differentiated phenotype characterized by reduced levels of MITF and increased levels of AXL and NGFR (Fig. 5E and Supplementary Fig. 8F).

Using the opposite strategy, we then asked whether ectopic expression of FSCN1 was able to revert the mesenchymal-like phenotype and restore drug sensitivity in BRAFi-resistant melanoma cells. Resistant cells transduced for stable FSCN1 overexpression displayed an increased proliferative rate compared to cells transduced with a control lentivirus (Fig. 5F). This effect was linked to diminished migratory abilities (Supplementary Fig. 9A). This phenotypic transition was further confirmed by Western Blot analysis of differentiation markers in various mesenchymal resistant cells, with increased expression of melanocytic markers (MITF, SOX10) and decreased levels of invasive markers (AXL, SOX9) as well as decreased production of ECM proteins and ECM-remodeling enzyme LOXL2 (Fig. 5G and Supplementary Fig. 9B). Finally, mirroring the effect of miR-143/-145 ASOs, forced expression of FSCN1 in M238R cells decreased viability in the presence of BRAFi (Fig. 5H). Overall these data underline the central function of the miR-143/-145/FSCN1 axis in the acquisition of an undifferentiated, mesenchymal-like cell state associated with therapy resistance.

### The miR-143-/145 cluster/FSCN1 axis regulates actin cytoskeleton dynamics and mechanopathways

Acquisition of the mesenchymal-like resistant state implies a massive cytoskeletal rearrangement reflected by morphological changes with cells assuming a flattened and spindle-like shape. Based on the key function of FSCN1 in F-actin microfilaments reorganization, we specifically analyzed the contribution of the miR-143/-145 cluster/FSCN1 axis on actin cytoskeleton dynamics. Transient overexpression of miR-143-3p and miR-145-5p reproduced these morphological changes, as shown by F-actin staining and increased cell area (Fig. 6A and Supplementary Fig. 10A). To better understand the crosstalk between ECM remodeling and rearranged actin dynamics, we performed immunofluorescent staining of focal adhesions, multi-protein structures that connect ECM to the acto-myosin cytoskeleton. An increased number of focal adhesions revealed by phospho-Paxillin staining characterized melanoma cells expressing miR-143-3p or miR-145-5p (Fig. 6B and Supplementary Fig. 10B). This result was also confirmed by Western Blot analysis of focal adhesion components such as phospho-FAK and phospho-SRC (Supplementary Fig. 10C). In addition, we observed an increase of phosphorylated and total forms of MLC2 and phosphorylated Signal Transducer and Activator of Transcription 3 (STAT3) upon cluster overexpression, suggesting the activation of the ROCK/JAK/STAT3 acto-myosin contractility pathway by the two miRNAs. We then investigated whether FSCN1 downregulation produced a similar effect on actin dynamics. Indeed, FSCN1 knockdown led to actin cytoskeleton reorganization with a significant cell area increase (Fig. 6C and Supplementary Fig. 10D) as well as an increased number of focal adhesions per cell (Fig. 6D and Supplementary Fig. 10E).

**Fig. 6.**
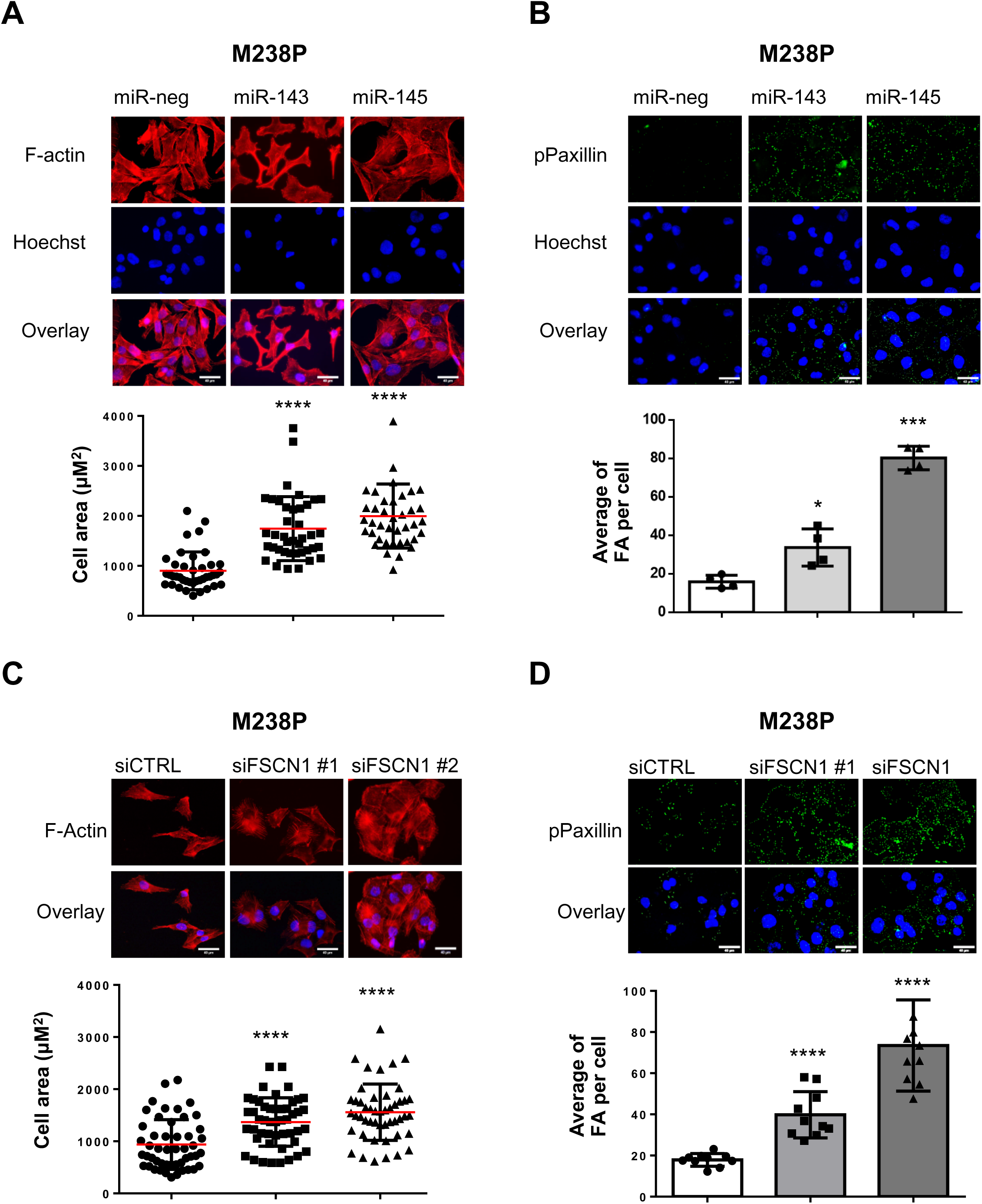
Regulation of actin cytoskeleton dynamics and focal adhesions by miR-143/-145 cluster /FSCN1 axis. (A-B) M238P cells were transfected with miR-143-3p, miR-145-5p or a control mimic (miR-neg) (72 h, 30 nM). **(A)** Quantification of cell area in cells stained for F-actin (red) and nuclei (blue). Data is represented as scatter plot with mean ± SD (n≥30 cells per condition). Mann-Whitney U test was used for statistical analysis. ****P≤0.0001. Scale bar 40 μM. **(B)** Quantification of focal adhesions number in cells stained for pPaxillin (green) and nuclei (blue). Focal adhesions number is represented as mean ± SD (n≥30 cells per condition). Each point represents the average number of focal adhesions per cell calculated for each field. Paired Student t-test has been used for statistical analysis. *P≤0.01 ***P≤0.001. FA, focal adhesion. Scale bar 40 μM. **(C-D)** M238P cells were transfected with two different sequences of siRNAs vs FSCN1 or with a control siRNA (72 h, 100 nM). **(C)** Quantification of cell area in cells stained for F-actin (red) and nuclei (blue). Data is represented as scatter plot with mean ± SD (n≥30 cells per condition). Mann-Whitney U test was used for statistical analysis. ****P≤0.0001. Scale bar 40 μM. **(D)** Quantification of focal adhesions number in cells stained for pPaxillin (green) and nuclei (blue). Focal adhesions number is represented as mean ± SD (n≥30 cells per condition). Each point represents the average number of focal adhesions per cell calculated for each field. Paired Student t-test has been used for statistical analysis. ****P≤0.0001. FA, focal adhesion. Scale bar 40 μM.

Acto-myosin remodeling critically regulates the cellular localization of mechanotransducers such as the Hippo pathway transcriptional co-activator YAP and the serum responsive factor co-activator MRTFA, two factors previously associated with resistance to MAPK-targeted therapies and pro-fibrotic responses (22, 23, 29, 33). Expression of miR-143-3p and miR-145-5p in therapy-naïve melanoma cells enhanced YAP and MRTFA nuclear localization as shown by immunofluorescent staining (Fig. 7A-B and Supplementary Fig. 11A-B). This increased YAP and MRTF activity was also confirmed by upregulated expression of several target genes (*CTGF*, *CYR61*, *AMOTL2*, *THBS1, AXL*), as shown by RT-qPCR analysis (Fig. 7C and Supplementary Fig. 11C). Again, these changes in cytoskeleton organization were reproduced by FSCN1 knockdown, with nuclear translocation of MRTFA and YAP (Fig. 7D-E and Supplementary Fig. 11D) and increased target gene expression (Fig. 7F). Finally, using the opposite strategy, we tested whether ectopic expression of FSCN1 was able to revert the constitutive activation of mechanical pathways typical of this cell state. Indeed, forced expression of FSCN1 in mesenchymal resistant cells significantly attenuated nuclear localization of YAP and MRTFA as well as their transcriptional activity (Supplementary Fig. 12A-C). Overall, our data underline the central function of the miR-143/-145/FSCN1 axis in the regulation of actin cytoskeleton dynamics and mechanopathways, leading to the acquisition of an undifferentiated, mesenchymal-like cell state associated with therapy resistance.

**Fig. 7.**
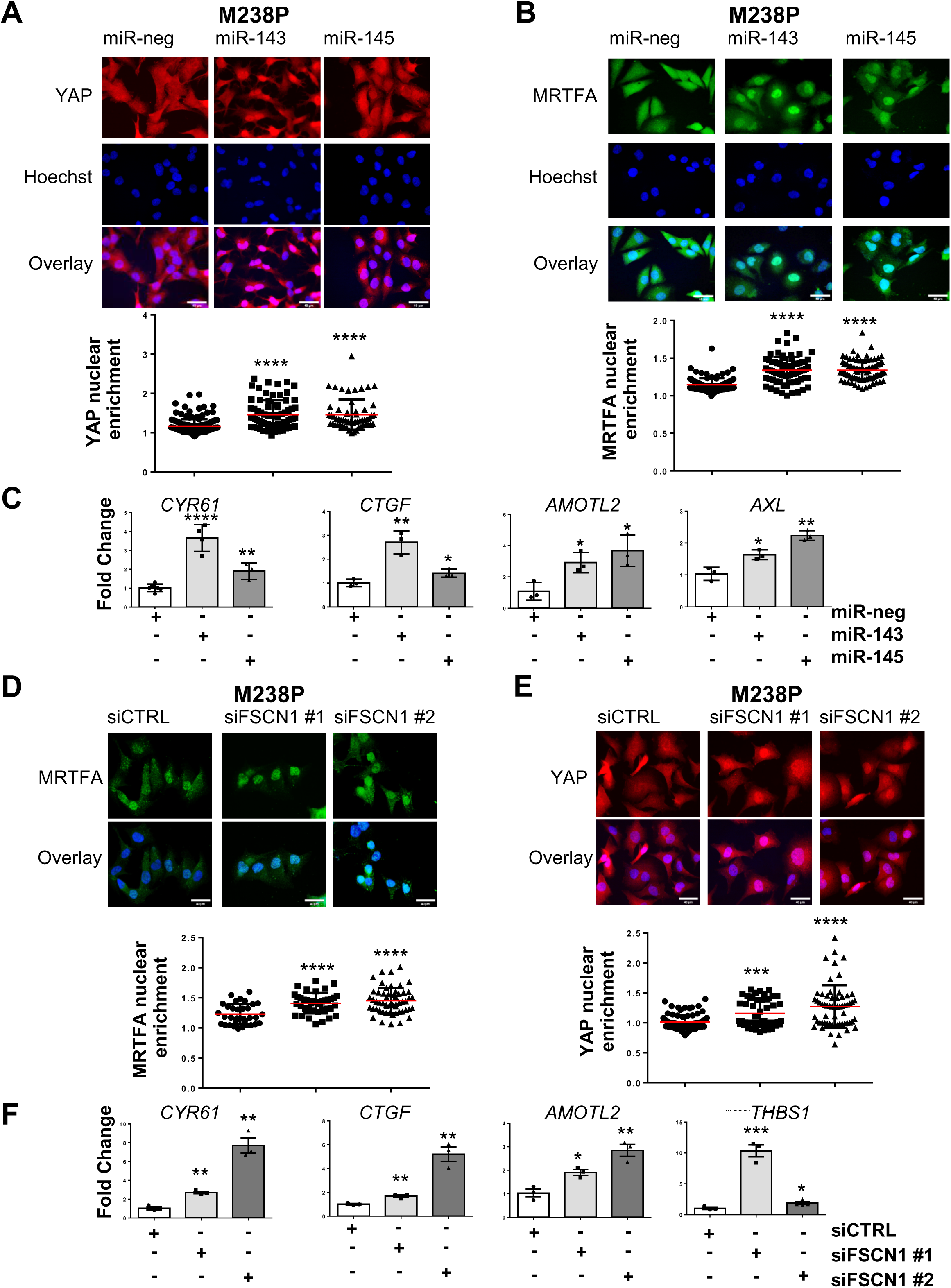
Regulation of mechanosensitive transcriptional coactivators YAP and MRTF by the miR-143/-145 cluster /FSCN1 axis. (A-C) M238P cells were transfected with miR-143-3p, miR-145-5p or a control mimic (miR-neg) (72 h, 30 nM). **(A-B)** Effect of miR-143-3p or miR-145-5p overexpression on YAP **(A)** and MRTFA **(B)** nuclear translocation by immunofluorescence. Data are represented as scatter plot with mean ± SD (n≥30 cells per condition). Mann-Whitney U test was used for statistical analysis. ****P≤0.0001. Scale bar 40 μM. **(C)** Effect of miR-143-3p or miR-145-5p overexpression on the expression of YAP/MRTF target genes assessed by RT-qPCR. Data are normalized to the expression in control cells. Data is represented as mean ± SD from a triplicate representative of at least 3 independent experiments. Paired Student t-test was used for statistical analysis. *P≤0.05, **P≤0.01, ****P≤0.0001. **(D-F)** M238P cells were transfected with two different sequences of siRNAs vs FSCN1 or with a control siRNA (72 h, 100 nM). **(D-E)** Effect of FSCN1 downregulation on MRTFA **(D)** and YAP1 **(E)** nuclear translocation assessed by immunofluorescence in M238P. Data are represented as scatter plot with mean ± SD (n≥30 cells per condition). Mann-Whitney U test was used for statistical analysis. ***P≤0.001, ****P≤0.0001. Scale bar 40 μM. **(F)** RT-qPCR analysis for the expression of MRTFA/YAP target genes in cells (M238P) transfected with the indicated siRNAs. Data are normalized to the expression in parental cells. Data is represented as mean ± SE from a triplicate representative of at least 3 independent experiments. Paired Student t-test has been used for statistical analysis. *P≤0.05, **P≤0.01, ***P≤0.001. Scale bar 40 μM.

## DISCUSSION

Treatments against advanced melanoma invariably end with therapy resistance and failure. Preventing resistance and tumor relapse on therapies targeting the MAPK oncogenic pathway still remains a challenge in successful melanoma clinical management. Our present study reveals that combination of the anti-fibrotic drug Nintedanib with targeted therapy provides therapeutic benefit in pre-clinical models of melanoma. We showed that Nintedanib is able to prevent the acquisition by melanoma cells of an undifferentiated mesenchymal-like phenotype, an aggressive cell state previously shown to be associated with the expression of pro-fibrotic markers, acquisition of myofibroblast/CAF-like activities and enhanced mechanosignaling as well as drug resistance (22, 23). Importantly, we provided evidence that the triplet combination BRAFi/MEKi/Nintedanib is active to normalize the fibrous collagen network, delay the onset of resistance and improve mice survival. We also confirmed the efficacy of this therapeutic combination in human BRAF^V600E^ mutant melanoma cells and described its potential to impair phenotype switching and improve response to MAPK targeted therapy (Fig. 8).

**Fig. 8.**
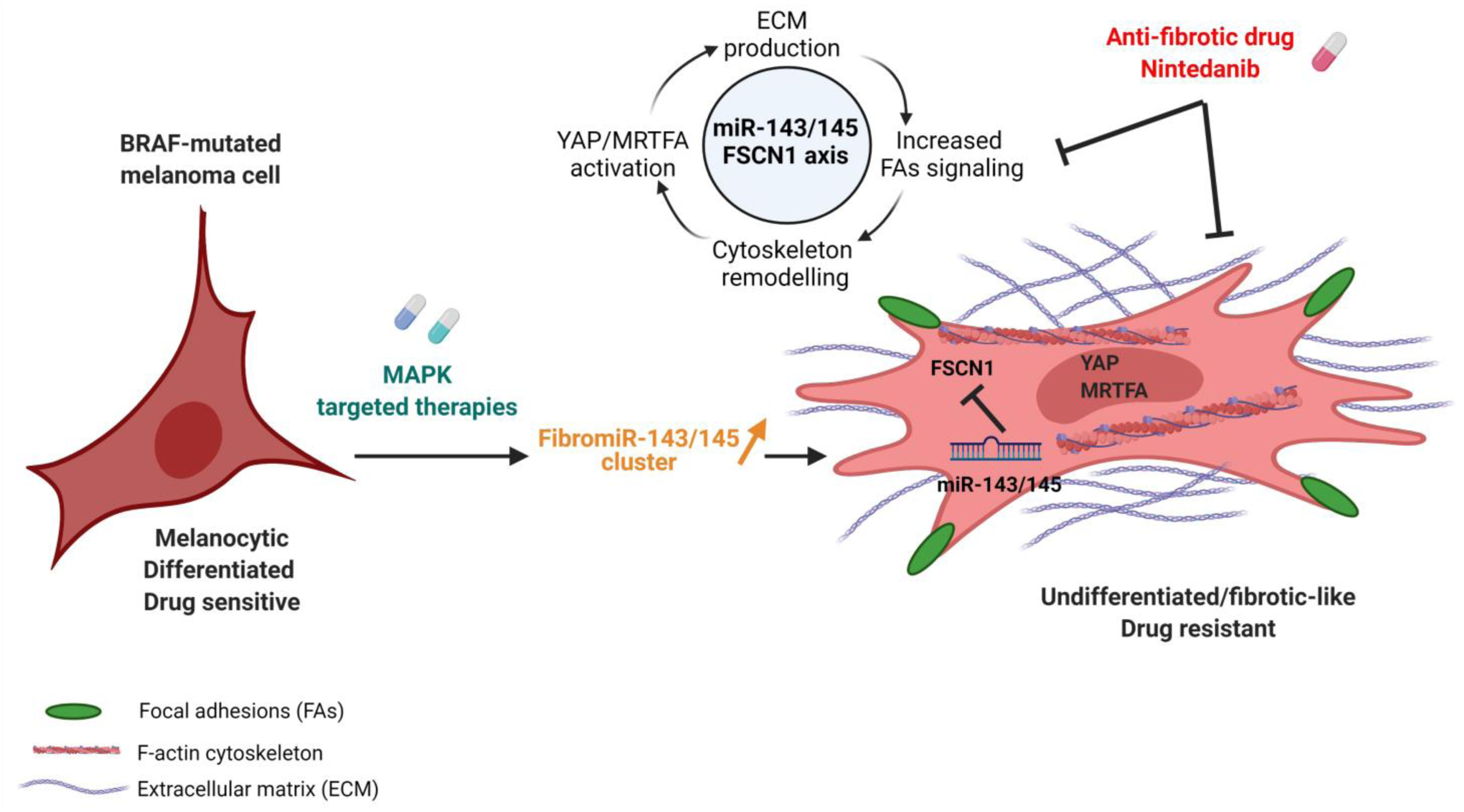
Proposed model for a role of the pro-fibrotic miR-143/-145 cluster in phenotypic plasticity-driven resistance induced by MAPK-targeted therapies and its potential targeting by Nintedanib. Created with <u>BioRender.com</u>

Nintedanib (BIBF-1120) is a multiple tyrosine kinase inhibitor, targeting PDGFR (α and β), FGFR-1, -2, -3, and -4 and VEGFR-1, -2, and -3 as well as several intracellular tyrosine kinases such as Src, Lck or Lyn. It has been approved for the treatment of Idiopathic Pulmonary Fibrosis (IPF) following several clinical trials demonstrating clinical efficacy in slowing disease progression (34). Nintedanib was shown to interfere with fundamental processes in lung fibrosis in a variety of *in vitro* assays performed on primary lung fibroblasts from patients with IPF, notably the inhibition of growth factor-induced proliferation/migration and TGF-β-induced myofibroblast activation as well as the down-regulation of ECM proteins (35). However, although substantial pre-clinical evidence demonstrates that Nintedanib has anti-fibrotic but also anti-inflammatory and anti-angiogenic activity, the exact contribution of inhibition of specific kinases to the activity of the drug in IPF has not been established and its precise anti-fibrotic mechanism(s) of action is not known.

In melanoma, the effects of Nintedanib are likely achieved through the normalization of the fibrotic and drug-protective ECM generated upon MAPK-targeted therapy exposure. We found that combined administration of Nintedanib and MAPK-targeted therapy dampens the increased miR-143/-145 cluster expression triggered by oncogenic BRAF pathway inhibition, suggesting that inhibition of ECM reprogramming in presence of Nintedanib is, at least partially, mediated by preventing upregulation of these two “FibromiRs”. Induction of the miR-143/-145 cluster paralleled the phenotypic switch associated with the undifferentiated mesenchymal-like phenotype and high expression levels of the two miRNAs are correlated with the mesenchymal MAPKi-resistant phenotype in all BRAF^V600E^ mutant human melanoma cell lines known to overexpress several RTKs including the PDGFR. Analysis of PDX models confirmed that expression levels of miR-143-3p and miR-145-5p are associated with a predominant invasive/undifferentiated transcriptomic profile in resistant lesions. Elevated levels of these miRNAs following BRAFi/MEKi treatment are likely due primarily to the direct inhibition of the MAPK pathway, as oncogenic signals including activation of the MAPK pathway strongly inhibit expression of the cluster in several epithelial cancers (36). In addition, we have shown a positive regulation of the cluster by PDGF or TGF-β signaling, as previously observed in the context of fibrosis and smooth muscle cell differentiation (37, 38). This observation supports the notion that pro-fibrotic signaling pathways typical of the mesenchymal resistance drive expression of the miR-143/-145 cluster in melanoma cells. Besides, the AKT pathway could also upregulate expression levels of the two miRNAs. Accordingly, previous studies stated that *PTEN* deletion favors the onset of a fibrotic phenotype in lung fibrosis and increased Fibronectin deposition in melanoma (20, 39). The observation that Nintedanib abrogated both AKT activation and miR-143/-145 expression in melanoma cells is in agreement with the importance of this pathway for acquisition and maintenance of drug resistance. Overall, our data indicate that Nintedanib can target both pro-fibrotic and survival pathways, mediated at least in part through PDGFR activation and converging to miR-143/-145 cluster expression.

The role of miR-143 and miR-145 in cancer has been widely debated in the last decade (40). The tumor suppressive role traditionally attributed to the cluster (41) has been challenged by recent genetic and cellular expression studies pointing mesenchymal cells as the main source of the cluster (42, 43). In melanoma, we disclosed that miR-143-3p and miR-145-5p promote the acquisition of an invasive and mesenchymal-like phenotype linked to drug adaptation and resistance. The importance of miR-143/145 cluster in the acquisition of this undifferentiated state is further highlighted with a loss-of-function approach showing that miR-143-3p and miR-145-5p inhibitors are able to limit ECM reprogramming and activation of mechanopathways, and improve anti-BRAF treatment efficacy. While further work using a combination of ASOs directed against the two mature miRNAs or the primary transcript is necessary to confirm these promising results in a melanoma xenograft model, we propose that the miR-143/-145 cluster may represent a novel attractive therapeutic target to prevent cells from switching to a mesenchymal/invasive state and tumor relapse after targeted therapy.

Our study shows that mechanistically the miR-143/-145 cluster functions in melanoma cells through targeting the cytoskeletal regulator FSCN1, one of the best hits identified by our screening, confirming previous studies indicating that FSCN1 is a direct target of both mature miRNAs (44, 45). FSCN1 has been widely studied in several malignancies for its role in promoting invasion and metastasis. However, a complete characterization of FSCN1 functions in melanoma is still missing and some published studies are controversial (46–48). Consistent with our study, FSCN1 downregulation was shown to inhibit melanoma cell proliferation (47) and to promote invasion (48). Interestingly, FSCN1 expression levels appear to be related to the differentiation stage of melanocytes and transient FSCN1 expression in melanoblasts precursors is required for their proliferation and migration, with FSCN1 knockout resulting in hypopigmentation in adult mice (47). Notably, miR-145-5p is also considered as a key regulator of the pigmentary process in melanocytes, a role mediated by the downregulation of pigmentation genes and melanosome trafficking components, including FSCN1 (49). These findings are in line with our data showing that FSCN1 downregulation drives phenotypic transition to an undifferentiated cell state associated with very low expression of the master regulator of melanocyte differentiation and function, MITF. FSCN1 downregulation may thus be exploited to generate lineage plasticity and revert to a poorly differentiated phenotype during drug adaptation of melanoma cells.

The miR-143/145 FSCN1 axis also directly modulates the dynamic crosstalk between the actin cytoskeleton and the ECM through the regulation of focal adhesion dynamics. This process is known to promote melanoma survival through FAK signaling and the ROCK pathway to induce acto-myosin-mediated contractile forces (50–52). The involvement of the miR-143/-145 cluster is also linked to a fine-tuning of mechanotransduction pathways. Enhanced YAP and MRTFA nuclear translocation reinforces the fibrotic-like phenotype promoted by the cluster and probably facilitates resistance acquisition, as previously demonstrated for these mechanotransducers (22, 29, 52). Interestingly, MRTFA has been involved in the transcriptional regulation of miR-143 and miR-145 expression (38, 53, 54), suggesting that this transcriptional state might be further stabilized by a positive feedback loop. Such regulatory loops between miRNAs and transcription factors have been previously described in the establishment and maintenance of melanoma phenotypic states (55, 56).

Despite the ability of FSCN1 downregulation to mimic the main functional effects observed by the ectopic expression of the miR-143/-145 cluster, we do not exclude the contribution of others targets in the acquisition of the mesenchymal resistant phenotype promoted by the cluster. FSCN1 knockdown failed to reproduce the global ECM signature reprogramming induced by the miR-143/-145 cluster. MiRNA target prediction tools identified a plethora of genes involved in cell cycle regulation, DNA damage response, inflammatory pathways, and actin-SRF regulatory network that need to be fully investigated in this context.

We conclude that our work opens new therapeutic avenues to prevent or delay the onset of targeted therapy resistance in melanoma. Our findings provide a rationale for designing clinical trials with Nintedanib and potentially other anti-fibrotic agents to enhance treatment efficacy in BRAF-mutated melanoma patients. We also bring an original mechanism of action directly linking the inhibition of the BRAF oncogenic pathway with the induction of the miR-143/-145 FibromiR cluster promoting the acquisition of a drug resistant, undifferentiated and mesenchymal-like cell state (Fig. 8). Finally, we propose the cluster as a new promising biomarker or druggable target to overcome non-genetic processes of phenotypic plasticity-driven therapeutic resistance.

## MATERIALS AND METHODS

### Cell lines and reagents

Isogenic pairs of Vemurafenib-sensitive and resistant cells (M229, M238, M249) were provided by R. Lo. UACC62 Vemurafenib-sensitive (UACC62P) and resistant cells (UACC62R) were provided by Neubig’s lab. 1205Lu cells were from Rockland. YUMM1.7 mouse melanoma cells were a kind gift from M. Bosenberg (24). Melanoma cells were cultured in Dulbecco’s modified Eagle medium (DMEM) supplemented with 7% FBS (Hyclone) and 1% penicillin/streptomycin. Resistant cells were continuously exposed to 1 μM of Vemurafenib. Cell lines were routinely tested for the absence of Mycoplasma by PCR.

Short-term cultures of patient melanoma cells MM034 and MM099 were generated in the laboratory of Pr G. Ghanem (ULB). Culture reagents were purchased from Thermo Fisher Scientific. BRAFi (PLX4032, Vemurafenib), MEKi (GSK1120212, Trametinib), SB431542, GSK690693, and Nintedanib (BIBF1120) were from Selleckem. Recombinant human TGF-β1 was from ImmunoTools. Recombinant human PDGF-BB was from Peprotech.

Information on all reagents used is provided in tables S4, S5 and S6.

### *In vivo* experiments

Mouse experiments were carried out according to the Institutional Animal Care and the local ethical committee (CIEPAL-Azur agreement NCE/2018-483). 4×10^5^ YUMM1.7 cells were injected in both flanks of C57BL/6 mice. Tumors were measured with caliper and treatments were started when the tumors reached a volume of 0.1 cm^3^, after randomization of mice into control and test groups. Vemurafenib (30 mg/kg), Trametinib (0.3 mg/kg), and Nintedanib (50 mg/kg) were administered by oral gavage three times per week. Control mice were treated with vehicle only. Animals were sacrificed when the tumors reached a volume of 1 cm^3^. After animal sacrifice, tumors were dissected, weighed and snap-frozen in liquid nitrogen for RNA or protein extraction and immunofluorescence analysis (embedded in OCT from Tissue-Tek). Tumors for picrosirius red staining were fixed in formalin. Melanoma cell-derived xenograft experiments performed on 6-week-old female athymic nude nu/nu mice were described in (22). Melanoma patient-derived xenografts models were established by TRACE (PDX platform; KU Leuven) using tissue from melanoma patients undergoing surgery at the University Hospitals KU Leuven. Written informed consent was obtained from all patients and all procedures were approved by the UZ Leuven Medical Ethical Committee (S54185/S57760/S59199) and carried out in accordance with the principles of the Declaration of Helsinki.

### Statistical analysis

Statistical analysis was performed using GraphPad Prism. Unpaired two-tailed Student’s T-test or unpaired two-tailed Mann Whitney test was used for statistical comparison between two groups. For comparisons between multiple groups, one-way ANOVA followed by Bonferroni’s *post*-*hoc* tests was used. For statistical analysis of cell confluence live imaging, two-way ANOVA was used. For statistical analysis of Kaplan-Meier curves, the log rank (Mantel-Cox) test was used. Results are given as mean ± SEM or mean ± SD

## Acknowledgments

We thank Roger Lo for M229P/R, M238P/R and M249P/R melanoma cells, Richard Neubig for UACC62 parental and resistant cell lines, Markus Bosenberg for YUMM1.7 cells and Ghanem Ghanem for short-term cultured melanoma cells. Melanoma PDX models are from the PDX facility TRACE (KULeuven, Belgium). We thank Patrick Brest and David Gilot for helpful discussions. We also thank the C3M animal room facility and the C3M imaging facility as well as the UCA GenomiX platform and the PVM Vectorology Platform.

## Author contributions

Conception and design of the work: SD, MD, BM, ST-D

Development of methodology: SD, AB, JF, ML, GV, MD, BM, ST-D

Acquisition of data: SD, AB, JF, ML, OM-B, CG, LL, CL, CM, MT, MC, AC, IB, FL, GV, J-CM

Analysis and interpretation of data: SD, AB, JF, MI, FL, MD, BM, ST-D

Writing – original draft: SD, BM, ST-D

Writing – review & editing: SD, CAG, GV, J-CM, MD, BM, ST-D

Administrative, technical or material support: MD, BM, ST-D

Study supervision: MD, BM, ST-D.

## Data and materials availability

Expression datasets that support the findings of this study have been deposited in the Gene Expression Omnibus SuperSerie record GSE171883 containing 3 distinct datasets under the following accession codes:

- Dataset 1: GSE171880. Effect of miR-143-3p or miR-145-5p mimics overexpression in M238P cells (microarrays).
- Dataset 2: GSE171881. RNA-Seq analysis of M238P stably expressing miR-143/-145 cluster.
- Dataset 3: GSE171882. Transcriptome analysis of M238R versus M238P using nanopore long reads sequencing.

All other data are available in the main text or in the supplementary materials.

## Supplementary Material and Methods

### Cell transduction

A DNA sequence containing the miR-143/145 cluster was cloned into a pLX307 vector by Sigma-Aldrich. The vector used for FSCN1 overexpression is described in [46]. Lentiviral particles were produced by the PVM Vectorology Platform in Montpellier, France. Melanoma cells were transduced as follows. After 20 min incubation of melanoma cells with lentiviral particles diluted in Optimem, complete medium (7% FBS) was added to the cells. Forty-eight hours after transduction, the process of antibiotic selection was started. For cells transduced for the miR-143/145 cluster overexpression, 1 μg/mL of puromycin was administered. For cells transduced for FSCN1 overexpression, 2 μg/mL of blasticidin was administered.

### RNAi studies

Non-targeting control and FSCN1 siRNA duplexes were designed by Sigma-Aldrich and used at a final concentration of 100 nM. Transfection was performed using Lipofectamine RNAiMAX (Life Technologies), according to the manufacturer’s instructions. Cells were analyzed 72 h post-transfection.

### miRNAs overexpression and inhibition

Pre-miRNAs -143-3p and -145-5p and control miRNA (miR-neg#1) were purchased from Ambion. LNA-based miRNAs inhibitors vs. miR-143-3p and miR-145-5p and the respective control (negative control A) were purchased from Qiagen. Pre-miRNAs were used at a final concentration of 30 nM, LNA inhibitors at a final concentration of 50 nM. Transfection was performed using Lipofectamine RNAiMAX (Life Technologies), according to the manufacturer’s instructions. Cells were analyzed 72 h post-transfection unless otherwise stated.

### Luciferase assay

Molecular constructs for luciferase assay were made in psiCHECK-2 vectors from Promega by cloning upstream of the Renilla luciferase gene annealed oligonucleotides based on the 3’UTR of target genes. HEK239 cells were plated on 96-well plates and co-transfected with 0.2 μg of psiCHECK-2 plasmid constructs and 10 nM of pre-miRNAs (miR-143-3p, miR-145-5p) or control pre-miRNA. Transfections were performed using Lipofectamine 3000, following the manufacturer’s instructions. Firefly and Renilla luciferase activities were measured using the Dual-Glo Luciferase assay kit by Promega 48 hours after transfection.

### Conditioned medium preparation

Medium conditioned by melanoma cells was harvested, centrifuged for 5 min at 2,500g and filtered with 0.22 μM filters to eliminate cell debris.

### Tumors and cells RNA extraction

Total RNA was extracted from tumors and cell samples with the miRNeasy Mini Kit (Qiagen) according to the manufacturer’s instructions.

### Real-time quantitative PCR

#### Gene expression

Protocol using the Step One (Applied Biosystem): 1 μg of extracted RNA was reverse transcribed into cDNA using the Multiscribe reverse transcriptase kit provided by Applied Biosystems. Primers were designed using PrimerBank or adopted from published studies. Gene expression levels were measured using Platinum SYBR Green qPCR Supermix (Fisher Scientific) and Step One thermocycler. Results from qPCR were normalized using the reference gene RPL32 and relative gene expression was quantified with the ΔΔCt method. Heatmaps describing gene expression fold changes were prepared using MeV software.

#### Protocol using the Biomark HD System Analysis (Fluidigm Corporation, USA)

cDNAs were prepared from 100 ng of RNA using Fluidigm Reverse Transcription Master Mix (Fluidigm PN 100-647297). Following a pre-amplification step (Fluidigm® PreAmp Master Mix and DELTAgene™ Assay kits) and exonuclease I treatment, samples diluted in Eva-Green® Supermix with Low ROX were loaded with primer reaction mixes in 96.96 Dynamic Array™ IFCs. Gene expression was then assessed on a Fluidigm BioMark HD instrument. Data were analyzed with real-time PCR analysis software (Fluidigm Corporation), and presented as relative gene expression according to the ΔΔCt method. Heatmaps depicting fold changes of gene expression were prepared using MeV software.

#### miRNAs expression

20 ng of extracted RNA was reverse transcribed into cDNA using the miRCURY LNA RT kit (Qiagen). Mature miRNAs expression levels were measured using the miRCURY LNA SYBR Green PCR kit (Qiagen). Results from qPCR were normalized using miR-16-5p and relative gene expression was quantified with the ΔΔCt method. miRCURY LNA miRNA PCR assays for detecting miR-143, miR-145, and miR-16 were purchased by Qiagen.

Information on primer sequences used in this study is provided in table S4 and S5.

### Immunoblot analysis and antibodies

Whole-cell lysates were prepared using RIPA buffer supplemented with protease and phosphatase inhibitor (Pierce, Fisher Scientific), briefly sonicated and centrifuged for 20 min, 4°C at 14000 rpm. Whole-cell lysates and conditioned media were separated using SDS-PAGE and transferred into PVDF membranes (GE Healthcare Life Science) for immunoblot analysis. Incubation of membranes with primary antibody was performed overnight. After washing, membranes were incubated with the peroxidase-conjugated secondary antibody. A chemiluminescence system (GE Healthcare Life Science) was used to develop blots. HSP60 or HSP90 were used as loading control. For immunoblot analysis of conditioned media experiments, Ponceau red staining was used as loading control.

Information on antibodies used in this study is provided in table S6.

### Immunofluorescence and microscopy

Cell monolayers were grown on glass coverslips or collagen-coated coverslips (10 μg/mL). After the indicated treatments, cells were washed in PBS, fixed in 4% PFA, permeabilized in PBS 0.3% Triton and blocked in PBS 5% goat serum. Coverslips were then incubated overnight at 4°C with primary antibody diluted in PBS 5% goat serum. Following 1 h incubation with Alexa-fluor conjugated secondary antibody, coverslips were mounted with Prolong antifade mounting reagent (ThermoFisher Scientific). Nuclei were stained with Hoechst 33342 (Life Technologies). F-actin was stained with Alexa Fluor 488 phalloidin (Fisher Scientific) or phalloidin-iFluor 594 (Abcam) reagents. Coverslips were imaged using a wide-field Leica DM5500B microscope.

### Fibrillar collagen imaging

Collagen in paraffin-embedded tumors was stained with picrosirius red using standard protocols. Tumor sections were analyzed by polarized light microscopy as described [26]. Images were acquired under polarized illumination using a light transmission microscope (Zeiss PALM, at 10x magnification). Fiber thickness was analyzed by the change in polarization color. Birefringence hue and amount were quantified as a percent of total tissue area using ImageJ software.

### Viability assay

After the indicated treatments, cells were stained with 0.04% crystal violet, 20% ethanol in PBS for 30 min. Following accurate washing of the plate, representative photographs were taken. The crystal violet dye was solubilized by 10% acetic acid in PBS and measured by absorbance at 595 nm.

### Proliferation assay

For real-time analysis of cell proliferation, 3×10^4^ cells were plated in complete medium in triplicates on 12-well plates. The Incucyte ZOOM imaging system (Essen Bioscience) was used. Phase-contrast pictures were taken every hour. Proliferation curves were generated using the IncuCyte cell proliferation assay software based on cell confluence.

### Cell cycle analysis

Cell cycle analysis was performed by flow cytometry analysis of cells stained with propidium iodide. After fixation in ice-cold 70% ethanol, cells were stained with 40 μg/mL propidium iodide in PBS with 100 μg/mL RNAse A. The samples were then analyzed on a BD FACSCanto cytometer.

### Migration and invasion assays

Migration properties of melanoma cells were tested using Boyden chambers containing polycarbonate membranes (8 μm pores transwell from Corning). After overnight starvation, 1×10^4^ cells were seeded on the upper side of the chambers placed on 24 well plates containing 10% FBS medium for 24 h, unless otherwise stated, at 37°C in 5% CO_2_. At the end of the experiment, cells migrated on the lower side of the chambers were fixed in 4% paraformaldehyde, stained for 15 min with Hoechst and imaged at the microscope (5 random fields per well). Nuclei counting was performed using the ImageJ software. To assess invasion properties of melanoma cells, transwells were coated with Matrigel (1 mg/mL) and cell solution was added on the top of the matrigel coating to simulate invasion through the extracellular matrix.

### Immunofluorescence analysis

Cell area was measured on cells stained for F-Actin using ImageJ. The nuclear/cytosolic ratio of YAP or MRTF was quantified by measuring the nuclear and cytosolic fluorescence intensity using ImageJ. The Hoechst staining was used to define nuclear versus cytosolic regions. Focal adhesions were quantified using ImageJ. Pictures were subjected to background subtraction (rolling: 10) before analysis, then “default threshold” was applied, followed by “analyze particles of object with a size 0.20 and infinity” to analyze the number of objects and their area. The number of focal adhesions was normalized to the total cell area.

### Microarray gene expression analysis

Total RNA integrity was tested with the Agilent BioAnalyser 2100 (Agilent Technologies). After labeling RNA samples with the Cy3 dye using the low RNA input QuickAmp kit (Agilent) following the manufacturer’s instruction, labeled cRNA probes were hybridized on 8×60K high-density SurePrint G3 gene expression human Agilent microarrays.

### RNA-sequencing

Short reads: Libraries were generated from 500ng of total RNAs using Truseq Stranded Total RNA kit (Illumina). Libraries were then quantified with KAPA library quantification kit (Kapa Biosystems) and pooled. 4nM of this pool were loaded on a high output flowcell and sequenced on an Illumina NextSeq500 sequencer using 2×75bp paired-end chemistry. Reads were aligned to the human genome release hg38 with STAR 2.5.2a as previously described [26].

Nanopore long reads: libraries were prepared according to the PCR-cDNA Barcoding protocol (SQK – PCB109). Briefly, 50 ng of total RNA was reverse-transcribed, barcoded and amplified by PCR (17 cycles) and sequencing adapters were added. The two barcoded libraries (CARMN RNA 238R and CARMN RNA 238S) were mixed 1:1, and 110 fmol was loaded on a PromethION flow cell (FLO-PRO002). Reads were processed with the FLAIR pipeline (https://doi.org/10.1038/s41467-020-15171-6). Raw reads were aligned to hg38 with minimap2 (version 2.17-r941). Misaligned splice sites were corrected according to the GENCODE v.35 annotations. High confidence isoforms were defined after grouping corrected reads of all samples sharing same unique splice junctions, by selecting for each group a representative isoform with confident TSS/TES and supported by more than 3 reads. Selected isoforms were quantified using minimap2 in each sample. Differential isoform expression and alternative splicing events significance were tested without replicates using ad-hoc scripts provided on the Brook’s lab Github (https://github.com/BrooksLabUCSC/FLAIR).

Statistical analysis and Biological Theme Analysis: Microarray data analyses were performed using R (http://www.r-project.org/). Quality control of expression arrays was performed using the Bioconductor package arrayQualityMetrics and custom R scripts. Additional analyses of expression arrays were performed using the Bioconductor package limma. Briefly, data were normalized using the quantile method. No background subtraction was performed. Replicated probes were averaged after normalization and control probes removed. Statistical significance was assessed using the limma moderated t-statistic Quality control of RNA-seq count data was assessed using in-house R scripts. Normalization and statistical analysis were performed using Bioconductor package DESeq2. All P-values were adjusted for multiple testing using the Benjamini-Hochberg procedure, which controls the false discovery rate (FDR). Differentially expressed genes were selected based on an adjusted p-value below 0.05. Enrichment in biological themes (Molecular function, Upstream regulators and canonical pathways) were performed using Ingenuity Pathway Analysis software (http://www.ingenuity.com/).

### miRNA targets analysis

MiRonTop is an online java web tool (http://www.genomique.info/) [31] integrating whole transcriptome expression data to investigate if specific miRNAs are involved in a specific biological system. MiRonTop classifies transcripts into two categories (’Upregulated’ and ’Downregulated’), based on thresholds for expression level, differential expression, and statistical significance. It then analyzes the number of predicted targets for each miRNA, according to the prediction software selected (Targetscan, exact seed search, TarBase).

## SUPPLEMENTARY TABLES

**Table S1.** List of themes corresponding to “Molecular functions” annotations associated with miR-143-3p and miR-145-5p mimics overexpression in human M238P melanoma cells identified by Ingenuity Pathway Analysis. M238P cells were transfected with miR-143, miR-145 or a control mimic (72 h, 30 nM). Expression profiles were determined with Agilent whole genome microarrays (Dataset 1). Z-scores calculated for each pathway (miR-143-3p or miR-145-5p vs miR-Neg) are indicated. Significant pathways are shown in progressively brighter shades of blue (repression) and orange (activation) according to their significance.

| Molecular Functions | - Log10 (P Val) |  |
| --- | --- | --- |
|  | miR-143-3p | miR-145-5p |
| Migration of cells | 12.13 | 15.13 |
| Cell movement | 11.08 | 14.26 |
| Cell proliferation of tumor cell lines | 11.80 | 11.09 |
| Invasion of cells | 10.24 | 8.80 |
| Migration of tumor cell lines | 10.46 | 7.65 |
| Cell movement of tumor cell lines | 8.93 | 9.02 |
| Organization of cytoplasm | 8.27 | 9.26 |
| Organization of cytoskeleton | 6.44 | 10.13 |
| Necrosis | 10.62 | 5.59 |
| Apoptosis | 10.07 | 5.72 |
| Cell death of tumor cell lines | 10.28 | 4.99 |
| Invasion of tumor cell lines | 9.31 | 5.86 |
| Cell survival | 8.75 | 5.23 |
| Cell viability | 8.90 | 4.96 |
| Microtubule dynamics | 4.54 | 9.04 |
| Cell cycle progression | 8.08 | 4.99 |
| Apoptosis of tumor cell lines | 6.70 | 4.56 |
| Cell viability of tumor cell lines | 6.02 | 4.61 |
| Phosphorylation of protein | 3.60 | 6.83 |
| Sprouting | 3.61 | 6.61 |
| Formation of cellular protrusions | 0.00 | 10.05 |
| Colony formation of cells | 6.12 | 3.70 |
| Biosynthesis of amide | 3.80 | 5.76 |
| Migration of cancer cells | 0.00 | 9.02 |
| Migration of tumor cells | 0.00 | 8.84 |
| Cell movement of tumor cells | 0.00 | 8.82 |
| Interphase | 8.15 | 0.00 |
| DNA replication | 7.64 | 0.00 |
| Transmigration of cells | 0.00 | 7.62 |
| Arrest in interphase | 6.86 | 0.00 |
| G1/S phase transition | 6.81 | 0.00 |
| G1 phase | 6.70 | 0.00 |
| Mitosis | 6.58 | 0.00 |
| Metabolism of DNA | 6.21 | 0.00 |
| Synthesis of DNA | 6.20 | 0.00 |
| Damage of chromosomes | 6.00 | 0.00 |
| Metabolism of reactive oxygen species | 5.44 | 0.00 |
| Chromosomal aberration | 5.37 | 0.00 |
| Entry into interphase | 5.30 | 0.00 |
| Chromosomal instability | 5.23 | 0.00 |
| Arrest in mitosis | 5.23 | 0.00 |
| Synthesis of glycosaminoglycan | 0.00 | 5.19 |
| Secretion of molecule | 0.00 | 5.16 |
| Synthesis of lipid | 0.00 | 5.16 |

**Table S2.**
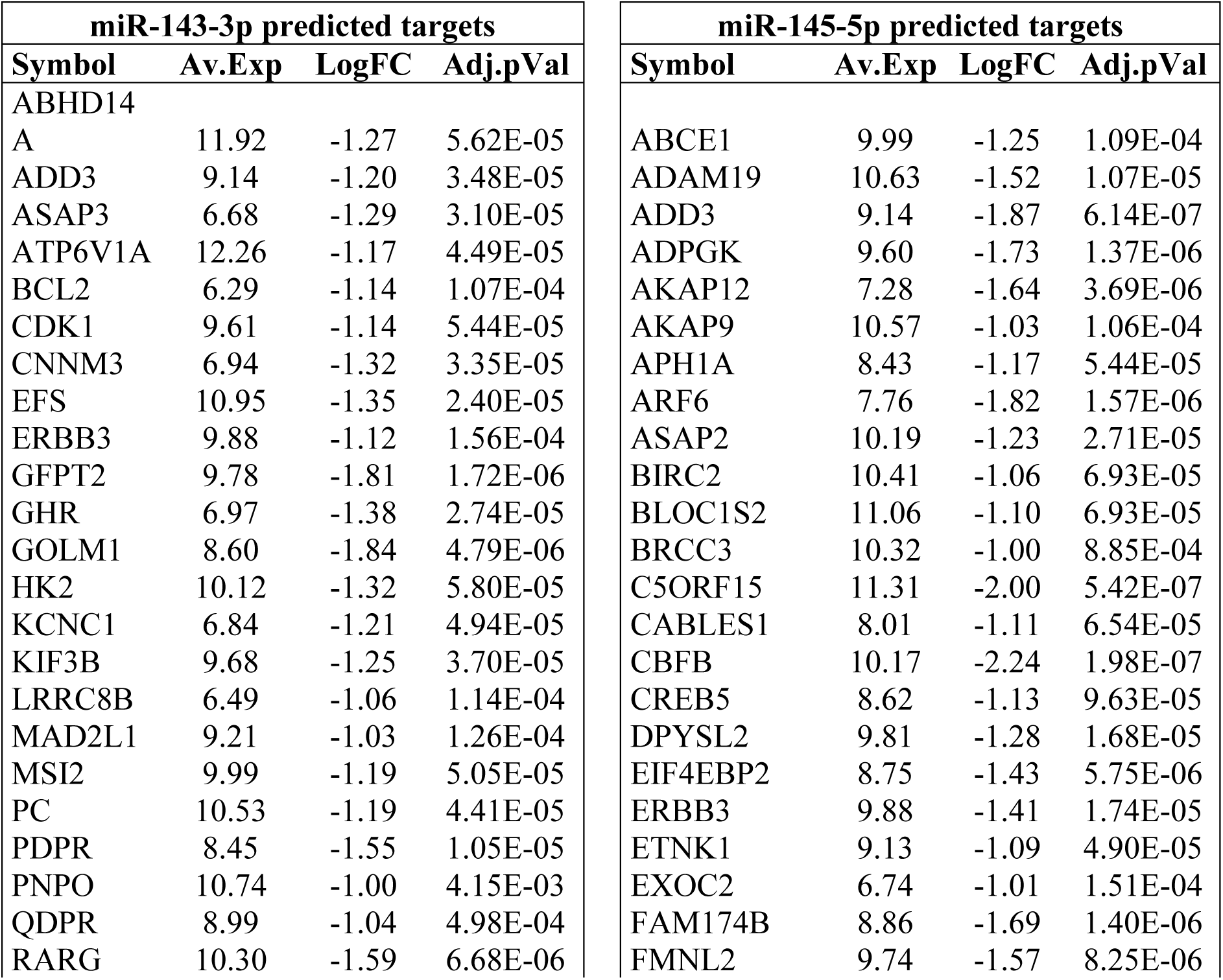
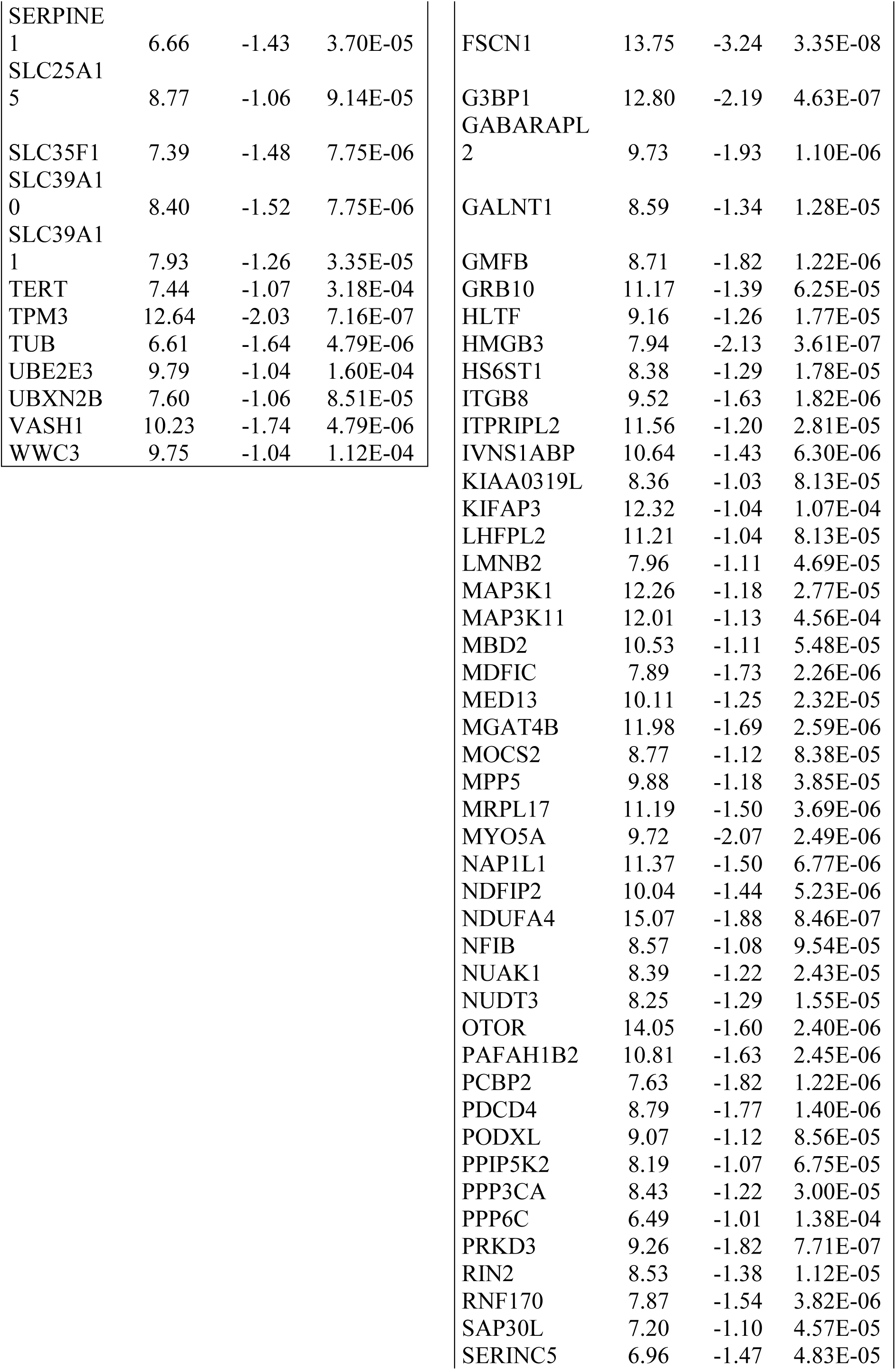
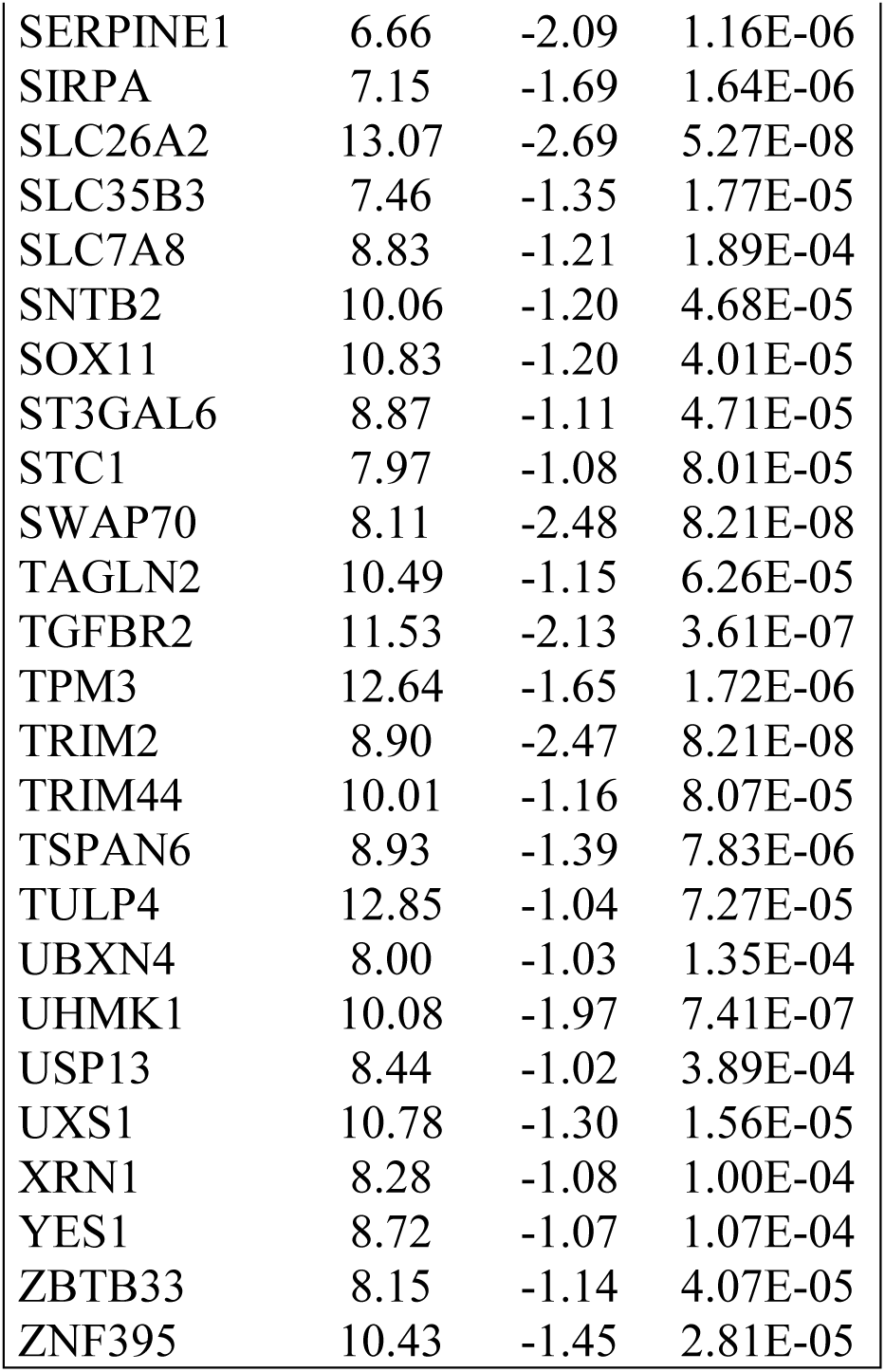
List of the main predicted targets for miR-143-3p and miR-145-5p significantly downregulated following mimics overexpression in human M238P melanoma cells using the bioinformatics tool miRonTop (https://www.genomique.info:8443/merge/index) [31]. M238P cells were transfected with miR-143, miR-145 or a control mimic (48 h, 30 nM). Expression profiles were determined with Agilent whole genome microarrays (Dataset 1). Predicted targets were obtained using the Targetscan database (conserved sites) (www.targetscan.org). Av.Exp: logarithm (base 2) of the average intensity ; logFC: logarithm (base 2) of the ratio of miR-143-3p or miR-145-5p vs miR-Neg; Adj.pVal: Benjamini-Hochberg adjusted pValue. The thresholds used for the analysis are: Av.Exp >6, LogFC<-1 and adj.pVal <0.05.

**Table S3.**
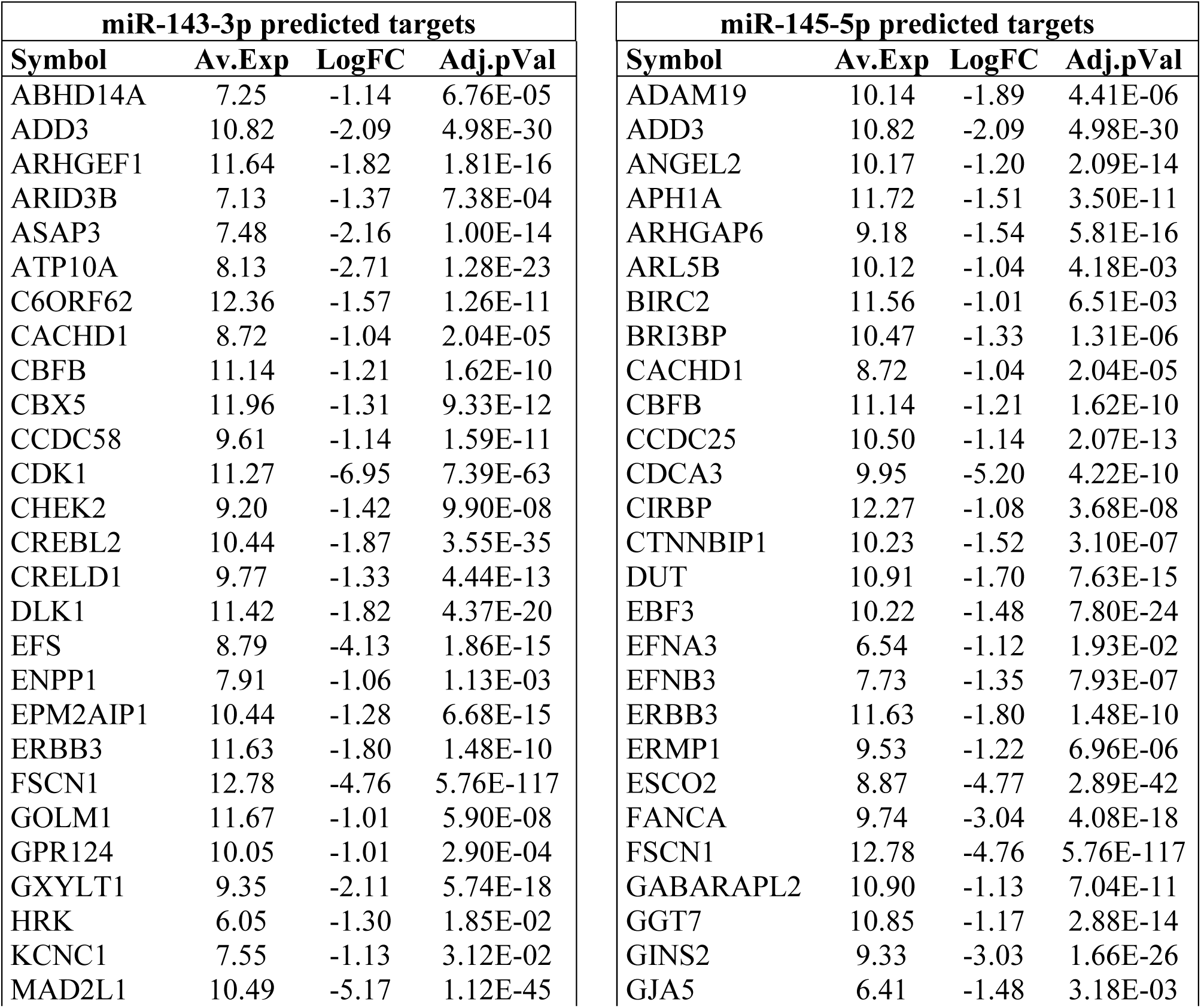
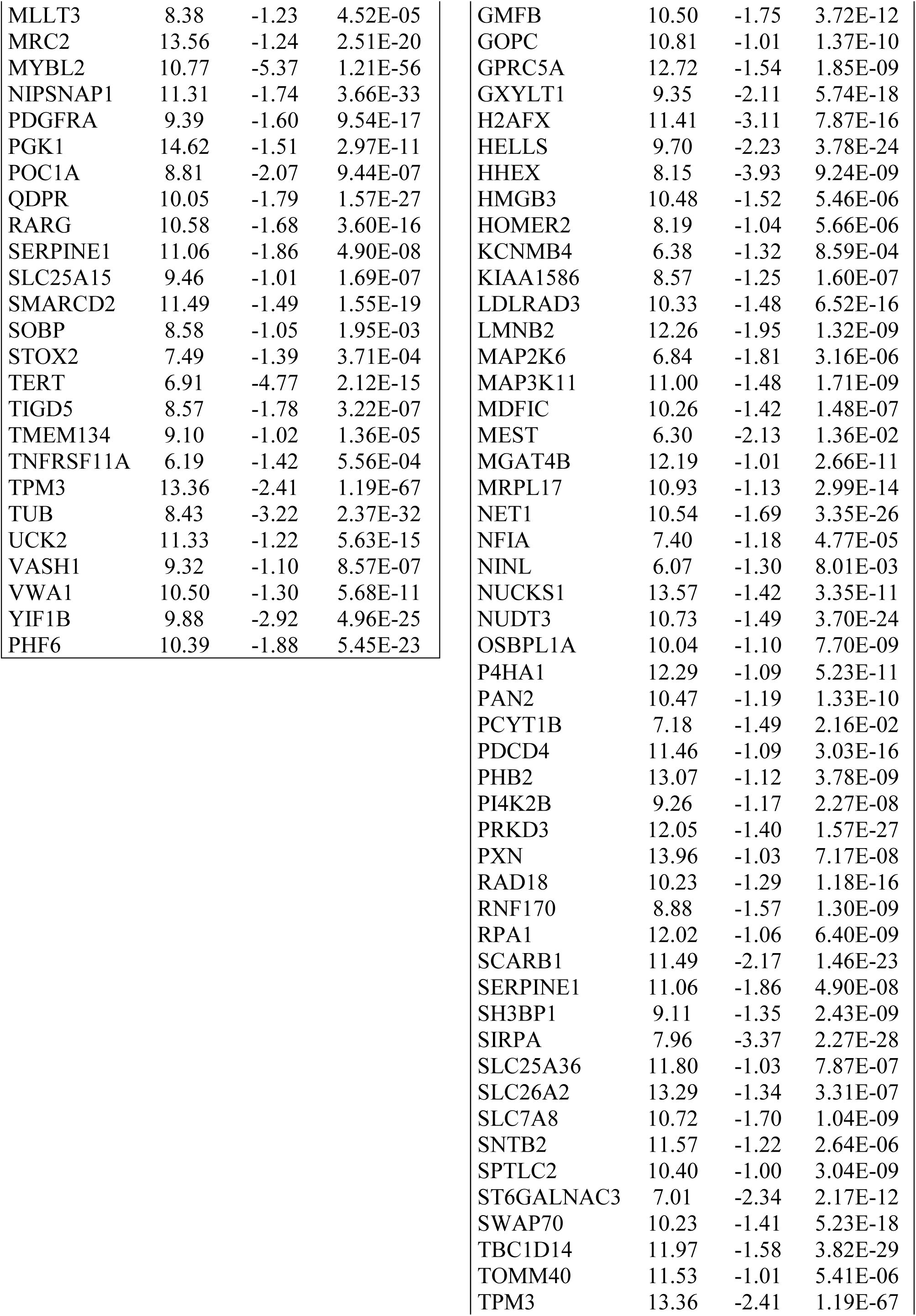
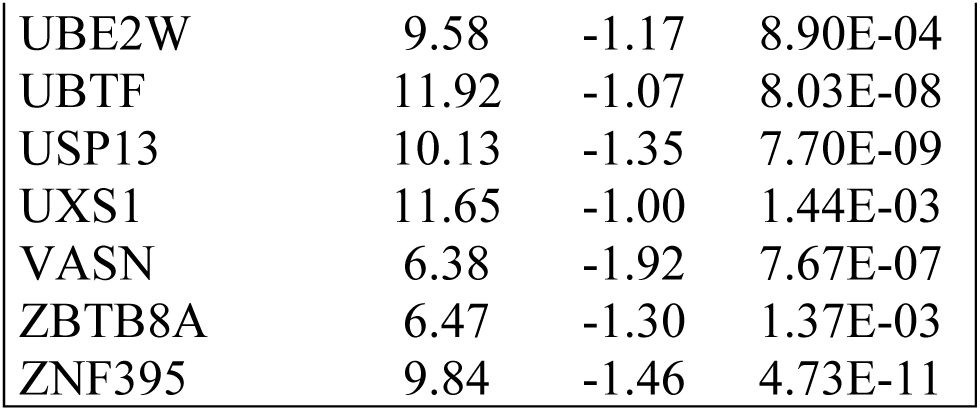
List of the main predicted targets for miR-143-3p and miR-145-5p significantly downregulated following stable overexpression of the miR-143/145 cluster in human M238P melanoma cells using the bioinformatics tool miRonTop (https://www.genomique.info:8443/merge/index) [31]. M238P melanoma cells were transduced with a lentivirus containing the sequence of the miR-143/145 cluster or a control vector. Expression profiles were determined with RNA-seq (Dataset 2). Predicted targets were obtained using the Targetscan database (conserved sites) (www.targetscan.org). Av.Exp: logarithm (base 2) of the base mean ; logFC: logarithm (base 2) of the ratio of miR-143/145 vs miR-Neg; Adj.pVal: Benjamini-Hochberg adjusted pValue. The thresholds used for the analysis are: Av.Exp >6, LogFC<-1 and adj.pVal <0.05.

| miR-143-3p predicted targets |  |  |  | miR-145-5p predicted targets |  |  |  |
| --- | --- | --- | --- | --- | --- | --- | --- |
| Symbol | Av.Exp | LogFC | Adj.pVal | Symbol | Av.Exp | LogFC | Adj.pVal |
| ABHD14A | 7.25 | -1.14 | 6.76E-05 | ADAM19 | 10.14 | -1.89 | 4.41E-06 |
| ADD3 | 10.82 | -2.09 | 4.98E-30 | ADD3 | 10.82 | -2.09 | 4.98E-30 |
| ARHGEF1 | 11.64 | -1.82 | 1.81E-16 | ANGEL2 | 10.17 | -1.20 | 2.09E-14 |
| ARID3B | 7.13 | -1.37 | 7.38E-04 | APH1A | 11.72 | -1.51 | 3.50E-11 |
| ASAP3 | 7.48 | -2.16 | 1.00E-14 | ARHGAP6 | 9.18 | -1.54 | 5.81E-16 |
| ATP10A | 8.13 | -2.71 | 1.28E-23 | ARL5B | 10.12 | -1.04 | 4.18E-03 |
| C6ORF62 | 12.36 | -1.57 | 1.26E-11 | BIRC2 | 11.56 | -1.01 | 6.51E-03 |
| CACHD1 | 8.72 | -1.04 | 2.04E-05 | BRI3BP | 10.47 | -1.33 | 1.31E-06 |
| CBFB | 11.14 | -1.21 | 1.62E-10 | CACHD1 | 8.72 | -1.04 | 2.04E-05 |
| CBX5 | 11.96 | -1.31 | 9.33E-12 | CBFB | 11.14 | -1.21 | 1.62E-10 |
| CCDC58 | 9.61 | -1.14 | 1.59E-11 | CCDC25 | 10.50 | -1.14 | 2.07E-13 |
| CDK1 | 11.27 | -6.95 | 7.39E-63 | CDCA3 | 9.95 | -5.20 | 4.22E-10 |
| CHEK2 | 9.20 | -1.42 | 9.90E-08 | CIRBP | 12.27 | -1.08 | 3.68E-08 |
| CREBL2 | 10.44 | -1.87 | 3.55E-35 | CTNNBIP1 | 10.23 | -1.52 | 3.10E-07 |
| CRELD1 | 9.77 | -1.33 | 4.44E-13 | DUT | 10.91 | -1.70 | 7.63E-15 |
| DLK1 | 11.42 | -1.82 | 4.37E-20 | EBF3 | 10.22 | -1.48 | 7.80E-24 |
| EFS | 8.79 | -4.13 | 1.86E-15 | EFNA3 | 6.54 | -1.12 | 1.93E-02 |
| ENPP1 | 7.91 | -1.06 | 1.13E-03 | EFNB3 | 7.73 | -1.35 | 7.93E-07 |
| EPM2AIP1 | 10.44 | -1.28 | 6.68E-15 | ERBB3 | 11.63 | -1.80 | 1.48E-10 |
| ERBB3 | 11.63 | -1.80 | 1.48E-10 | ERMP1 | 9.53 | -1.22 | 6.96E-06 |
| FSCN1 | 12.78 | -4.76 | 5.76E-117 | ESCO2 | 8.87 | -4.77 | 2.89E-42 |
| GOLM1 | 11.67 | -1.01 | 5.90E-08 | FANCA | 9.74 | -3.04 | 4.08E-18 |
| GPR124 | 10.05 | -1.01 | 2.90E-04 | FSCN1 | 12.78 | -4.76 | 5.76E-117 |
| GXYLT1 | 9.35 | -2.11 | 5.74E-18 | GABARAPL2 | 10.90 | -1.13 | 7.04E-11 |
| HRK | 6.05 | -1.30 | 1.85E-02 | GGT7 | 10.85 | -1.17 | 2.88E-14 |
| KCNC1 | 7.55 | -1.13 | 3.12E-02 | GINS2 | 9.33 | -3.03 | 1.66E-26 |
| MAD2L1 | 10.49 | -5.17 | 1.12E-45 | GJA5 | 6.41 | -1.48 | 3.18E-03 |
| MLLT3 | 8.38 | -1.23 | 4.52E-05 | GMFB | 10.50 | -1.75 | 3.72E-12 |
| MRC2 | 13.56 | -1.24 | 2.51E-20 | GOPC | 10.81 | -1.01 | 1.37E-10 |
| MYBL2 | 10.77 | -5.37 | 1.21E-56 | GPRC5A | 12.72 | -1.54 | 1.85E-09 |
| NIPSNAP1 | 11.31 | -1.74 | 3.66E-33 | GXYLT1 | 9.35 | -2.11 | 5.74E-18 |
| PDGFRA | 9.39 | -1.60 | 9.54E-17 | H2AFX | 11.41 | -3.11 | 7.87E-16 |
| PGK1 | 14.62 | -1.51 | 2.97E-11 | HELLS | 9.70 | -2.23 | 3.78E-24 |
| POC1A | 8.81 | -2.07 | 9.44E-07 | HHEX | 8.15 | -3.93 | 9.24E-09 |
| QDPR | 10.05 | -1.79 | 1.57E-27 | HMGB3 | 10.48 | -1.52 | 5.46E-06 |
| RARG | 10.58 | -1.68 | 3.60E-16 | HOMER2 | 8.19 | -1.04 | 5.66E-06 |
| SERPINE1 | 11.06 | -1.86 | 4.90E-08 | KCNMB4 | 6.38 | -1.32 | 8.59E-04 |
| SLC25A15 | 9.46 | -1.01 | 1.69E-07 | KIAA1586 | 8.57 | -1.25 | 1.60E-07 |
| SMARCD2 | 11.49 | -1.49 | 1.55E-19 | LDLRAD3 | 10.33 | -1.48 | 6.52E-16 |
| SOBP | 8.58 | -1.05 | 1.95E-03 | LMNB2 | 12.26 | -1.95 | 1.32E-09 |
| STOX2 | 7.49 | -1.39 | 3.71E-04 | MAP2K6 | 6.84 | -1.81 | 3.16E-06 |
| TERT | 6.91 | -4.77 | 2.12E-15 | MAP3K11 | 11.00 | -1.48 | 1.71E-09 |
| TIGD5 | 8.57 | -1.78 | 3.22E-07 | MDFIC | 10.26 | -1.42 | 1.48E-07 |
| TMEM134 | 9.10 | -1.02 | 1.36E-05 | MEST | 6.30 | -2.13 | 1.36E-02 |
| TNFRSF11A | 6.19 | -1.42 | 5.56E-04 | MGAT4B | 12.19 | -1.01 | 2.66E-11 |
| TPM3 | 13.36 | -2.41 | 1.19E-67 | MRPL17 | 10.93 | -1.13 | 2.99E-14 |
| TUB | 8.43 | -3.22 | 2.37E-32 | NET1 | 10.54 | -1.69 | 3.35E-26 |
| UCK2 | 11.33 | -1.22 | 5.63E-15 | NFIA | 7.40 | -1.18 | 4.77E-05 |
| VASH1 | 9.32 | -1.10 | 8.57E-07 | NINL | 6.07 | -1.30 | 8.01E-03 |
| VWA1 | 10.50 | -1.30 | 5.68E-11 | NUCKS1 | 13.57 | -1.42 | 3.35E-11 |
| YIF1B | 9.88 | -2.92 | 4.96E-25 | NUDT3 | 10.73 | -1.49 | 3.70E-24 |
| PHF6 | 10.39 | -1.88 | 5.45E-23 | OSBPL1A | 10.04 | -1.10 | 7.70E-09 |
|  |  |  |  | P4HA1 | 12.29 | -1.09 | 5.23E-11 |
|  |  |  |  | PAN2 | 10.47 | -1.19 | 1.33E-10 |
|  |  |  |  | PCYT1B | 7.18 | -1.49 | 2.16E-02 |
|  |  |  |  | PDCD4 | 11.46 | -1.09 | 3.03E-16 |
|  |  |  |  | PHB2 | 13.07 | -1.12 | 3.78E-09 |
|  |  |  |  | PI4K2B | 9.26 | -1.17 | 2.27E-08 |
|  |  |  |  | PRKD3 | 12.05 | -1.40 | 1.57E-27 |
|  |  |  |  | PXN | 13.96 | -1.03 | 7.17E-08 |
|  |  |  |  | RAD18 | 10.23 | -1.29 | 1.18E-16 |
|  |  |  |  | RNF170 | 8.88 | -1.57 | 1.30E-09 |
|  |  |  |  | RPA1 | 12.02 | -1.06 | 6.40E-09 |
|  |  |  |  | SCARB1 | 11.49 | -2.17 | 1.46E-23 |
|  |  |  |  | SERPINE1 | 11.06 | -1.86 | 4.90E-08 |
|  |  |  |  | SH3BP1 | 9.11 | -1.35 | 2.43E-09 |
|  |  |  |  | SIRPA | 7.96 | -3.37 | 2.27E-28 |
|  |  |  |  | SLC25A36 | 11.80 | -1.03 | 7.87E-07 |
|  |  |  |  | SLC26A2 | 13.29 | -1.34 | 3.31E-07 |
|  |  |  |  | SLC7A8 | 10.72 | -1.70 | 1.04E-09 |
|  |  |  |  | SNTB2 | 11.57 | -1.22 | 2.64E-06 |
|  |  |  |  | SPTLC2 | 10.40 | -1.00 | 3.04E-09 |
|  |  |  |  | ST6GALNAC3 | 7.01 | -2.34 | 2.17E-12 |
|  |  |  |  | SWAP70 | 10.23 | -1.41 | 5.23E-18 |
|  |  |  |  | TBC1D14 | 11.97 | -1.58 | 3.82E-29 |
|  |  |  |  | TOMM40 | 11.53 | -1.01 | 5.41E-06 |
|  |  |  |  | TPM3 | 13.36 | -2.41 | 1.19E-67 |

|  |  |  |  |
| --- | --- | --- | --- |
| UBE2W | 9.58 | -1.17 | 8.90E-04 |
| UBTF | 11.92 | -1.07 | 8.03E-08 |
| USP13 | 10.13 | -1.35 | 7.70E-09 |
| UXS1 | 11.65 | -1.00 | 1.44E-03 |
| VASN | 6.38 | -1.92 | 7.67E-07 |
| ZBTB8A | 6.47 | -1.30 | 1.37E-03 |
| ZNF395 | 9.84 | -1.46 | 4.73E-11 |

**Table S4.**
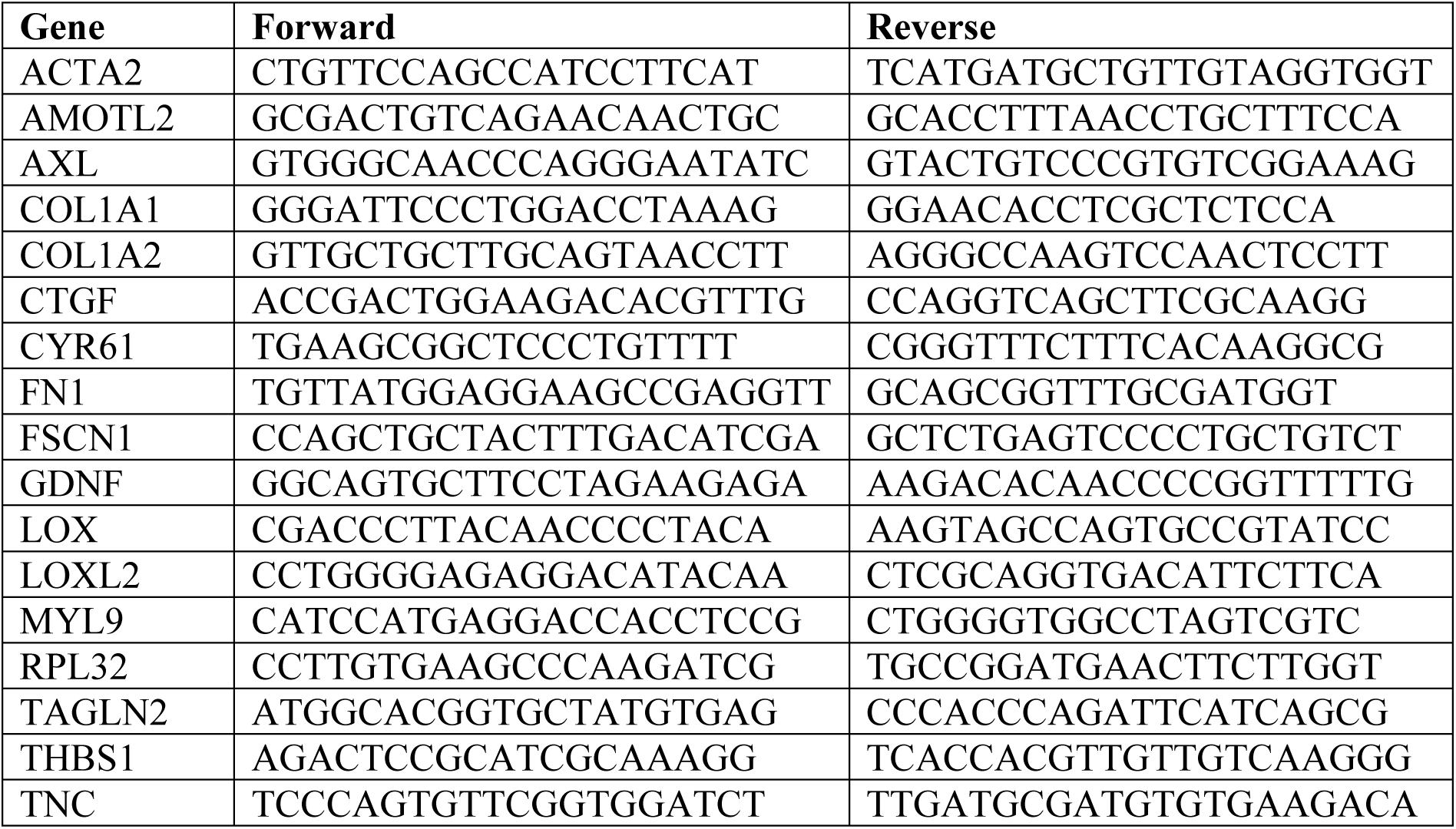
Human primers sequences used in the study.

**Table S5.**
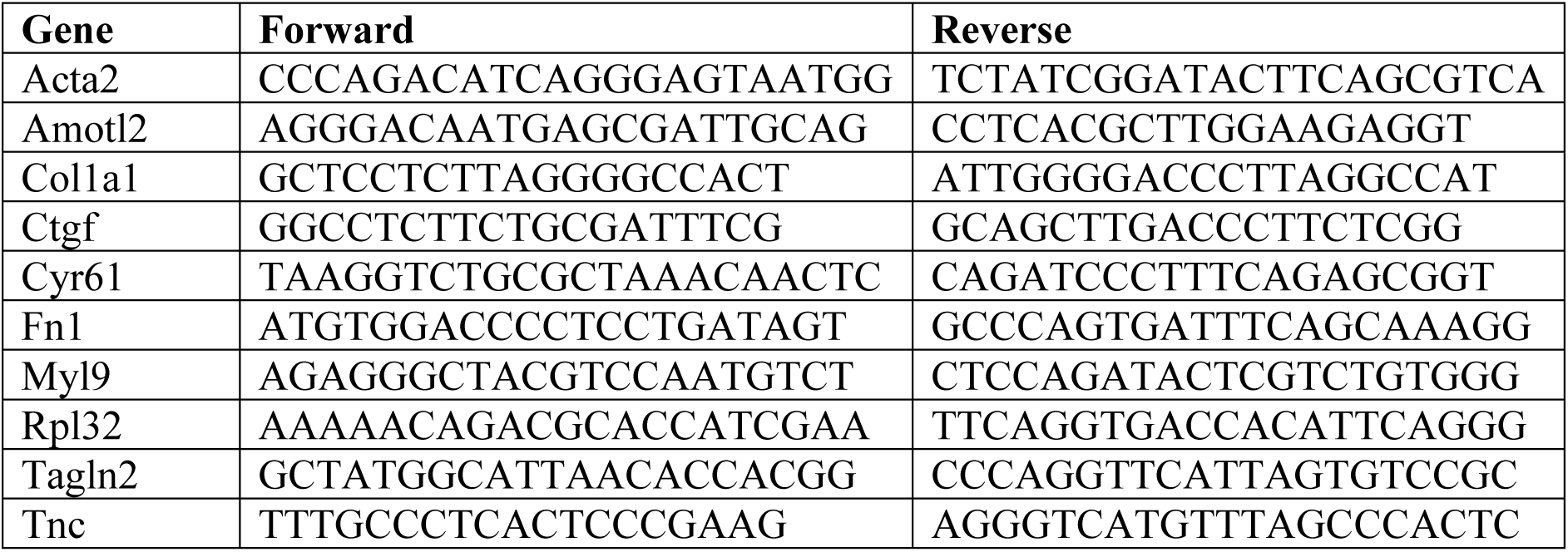
Mouse primers sequences used in the study.

**Table S6.** List of antibodies used in the study.

| **Primary Antibody** | **Company** | **Catalog Number** | **Dilution** |
| --- | --- | --- | --- |
| p-AKT (Ser473) | Cell Signaling | 9271 | WB 1:1000 |
| AKT (pan) | Cell Signaling | 4691 | WB 1:1000 |
| αSMA | Abcam | ab5694 | WB 1:1000 |
| AXL | Cell Signaling | 4566 | WB 1:1000 |
| CDK2 | Santa Cruz | sc-6248 | WB 1:500 |
| COL1 | Abcam | ab34710 | WB 1:3000 |
| CCND1 | BD Biosciences | 556470 | WB 1:1000 |
| E2F1 | Cell Signaling | 3742 | WB 1:1000 |
| EGFR | Santa Cruz | sc-373746 | WB 1:500 |
| p-ERK1/2<br>(Thr202/Tyr204) | Cell Signaling | 9101 | WB 1:1000 |
| ERK2 | Santa Cruz | sc-1647 | WB 1:500 |
| FAK | Upstate | 05-182 | WB 1:1000 |
| p-FAK (Tyr397) | Cell Signaling | 3283 | WB 1:500 |
| FN1 | Santa Cruz | sc-8422 | WB 1:500, IF 1:100 |
| FSCN1 | Santa Cruz | sc-21743 | IF 1:100 |
| FSCN1 | Proteintech | 66321-1-Ig | WB 1:1000 |
| HSP60 | Santa Cruz | sc-57840 | WB 1:500 |
| HSP90 | Santa Cruz | sc-13119 | WB 1:500 |
| LOX | Novus Biologicals | NB100-2527SS | WB 1:1000 |
| LOXL2 | R&D Systems | AF2639 | WB 1:1000 |
| MITF | Sigma | HPA003259 | WB 1:1000 |
| MLC2 | Cell Signaling | 3672 | WB 1:1000 |
| p-MLC2<br>(Thr18/Ser19) | Cell Signaling | 3674 | WB 1:500 |
| MRTFA | Santa Cruz | sc-390324 | IF 1:100 |
| NGFR (p75NTR) | Cell Signaling | 8238 | WB 1:1000 |
| p27 Kip1 | Cell Signaling | 3686 | WB 1:1000 |
| Paxillin | BD Biosciences | P13520 | WB 1:3000 |
| p-Paxillin (Tyr118) | Cell Signaling | 2541 | WB 1:1000 IF 1:50 |
| PDGFRβ | Santa Cruz | sc-374573 | WB 1:500 |
| p-Rb | Cell Signaling | 9308 | WB 1:1000 |
| Rb | Cell Signaling | 9309 | WB 1:1000 |
| p-SMAD3<br>(Ser433/435)/SMAD1<br>(Ser463/465) | Cell Signaling | 9514 | WB 1:1000 |
| SMAD1/2/3 | Santa Cruz | sc-7960 | WB 1:500 |
| SOX9 | Santa Cruz | sc-166505 | WB 1:500 |
| SOX10 | Cell Signaling | 89356 | WB 1:1000 |
| pSrc family (Tyr416) | Cell Signaling | 6943 | WB 1:1000 |
| Src | Cell Signaling | 2109 | WB 1:1000 |
| STAT3 | Cell Signaling | 9139 | WB 1:1000 |
| p-STAT3 (Tyr705) | Cell Signaling | 9145 | WB 1:1000 |
| Survivin | Cell Signaling | 2808 | WB 1:1000 |
| TAGLN2 | Genetex | GTX115082 | WB 1:1000 |
| THBS1 | Santa Cruz | sc-393504 | WB 1:500 |
| TNC | R&D Systems | AF3358 | WB 1:1000 |
| YAP | Cell Signaling | 14074 | IF 1:200 |

| **Secondary Antibody** | **Company** | **Catalog Number** | **Dilutions** |
| --- | --- | --- | --- |
| Anti-mouse IgG, HRP-linked antibody | Cell Signaling | 7076 | WB 1:2000 |
| Anti-rabbit IgG, HRP-linked antibody | Cell Signaling | 7074 | WB 1:2000 |
| Anti-goat IgG, HRP-linked antibody | Santa Cruz | sc-2354 | WB 1:5000 |
| Goat- anti-mouse, Alexa Fluor® 488 | Invitrogen | A11001 | IF 1:200 |
| Goat anti-mouse, Alexa Fluor® 594 | Invitrogen | A11005 | IF 1:200 |
| Goat anti-rabbit, Alexa Fluor® 594 | Invitrogen | A11012 | IF 1:200 |

## SUPPLEMENTARY FIGURE LEGENDS

**Fig. S1.**
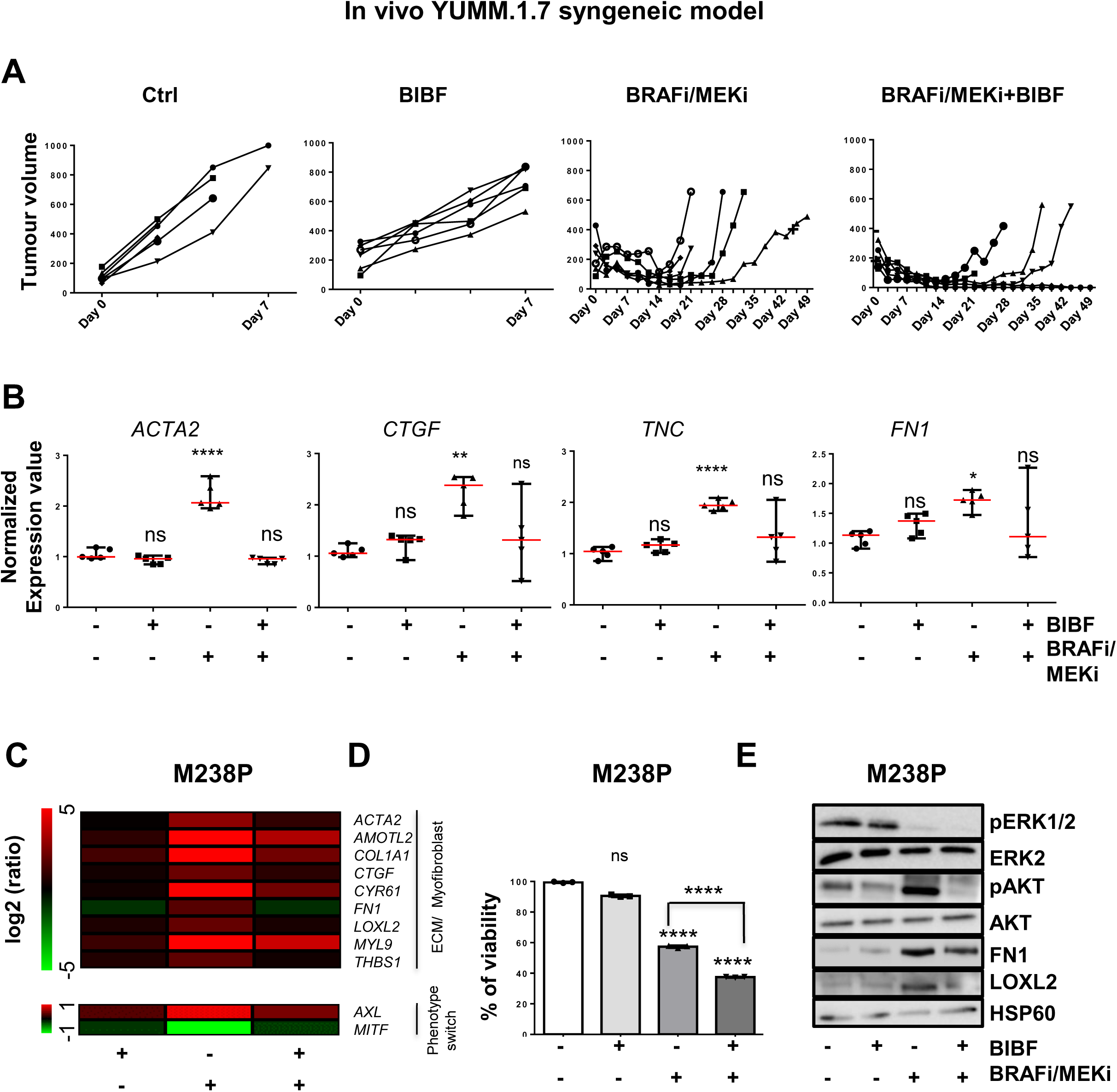
Administration of Nintedanib/BIBF1120 re-sensitizes melanoma cells to MAPK targeted therapies, delays tumor relapse and normalizes MAPKi-induced ECM remodeling and miR-143/145 expression. (A-D) YUMM1.7 cells were subcutaneously inoculated into C57BL/6 mice and when tumors reached 100 mm^3^ mice were treated with the indicated therapies. (**A**) Individual graphics showing tumor growth following treatment. **(B)** Normalized expression of myofibroblast/CAF and ECM-related genes assessed by RT-qPCR in individual tumors treated as indicated. Data is represented as median with range (n=5). One-way ANOVA has been used for statistical analysis. *P≤0.05, **P≤0.01, ****P≤0.0001. **(C-E)** Human M238P cells were treated with MAPK-targeted therapies (BRAFi, Vemurafenib **+** MEKi, Trametinib) (1μM), BIBF1120 (BIBF) (2 µM) or with MAPK-targeted therapies (1 µM) plus BIBF (2 µM) for 72 h. **(C)** Heatmap showing the expression of ECM, myofibroblast/CAF markers and phenotype switch markers by RT-qPCR (n=3). **(D)** Crystal violet viability assay of M238P cells treated with MAPK-targeted therapies (BRAFi, Vemurafenib **+** MEKi, Trametinib) (1μM), BIBF1120 (BIBF) (2 µM) or with MAPK-targeted therapies (1 µM) plus BIBF (2 µM) for 72 h. Paired Student t-test was used for statistical analysis. ****P≤0.0001. Data is represented as mean ± SD from a triplicate representative of 3 independent experiments. **(E)** Western blot showing the expression of ECM, myofibroblast/CAF markers and activation levels of signaling pathways (AKT and MAPK) in the different conditions.

**Fig. S2.**
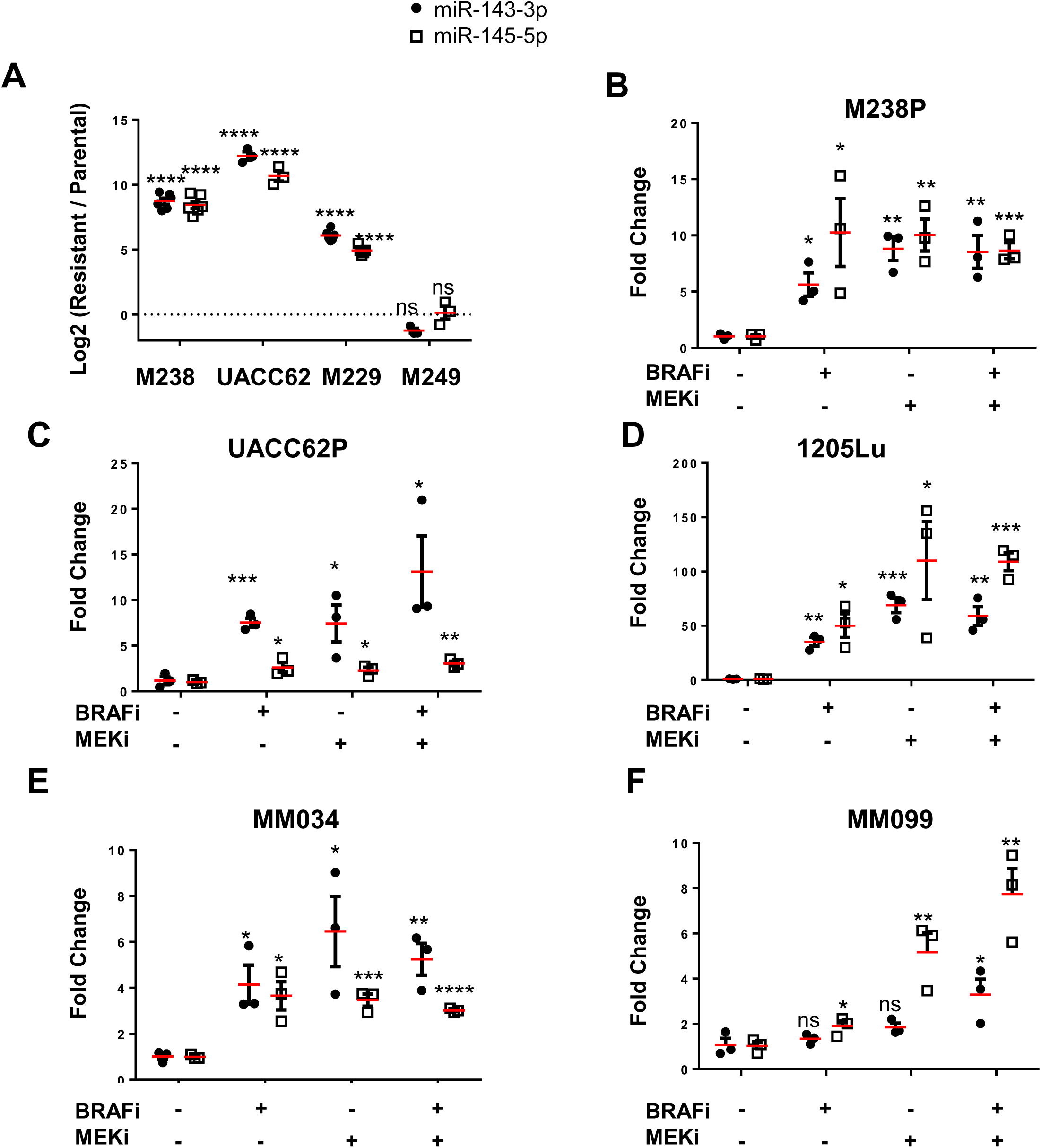
High expression of miR-143/145 is correlated with an undifferentiated/mesenchymal-like BRAFi-resistant phenotype in melanoma cells. **(A)** Relative miRNA expression levels have been quantified in parental (P) and paired resistant (R) cells (M238, UACC62, M229, M249) by RT-qPCR. Log2 (R/P) is shown for each cell line. **(B-F)** Relative miRNA expression levels have been quantified in human melanoma cell lines (M238P, UACC62P, 1205Lu) or short term patient-derived cell lines (MM034, MM099) treated or not for 72 h with MAPK-targeted therapies (BRAFi, Vemurafenib 3 µM), (MEKi, Trametinib 1 µM), (BRAFi plus MEKi, 1 µM) by RT-qPCR and normalized to miR-16-5p. **(A-F)** Data is represented as mean ± SE from a triplicate representative of at least 3 independent experiments. Paired Student t-test has been used for statistical analysis. *P≤0.05 **P≤0.01 ***P≤0.001 ****P≤0.0001.

**Fig. S3.**
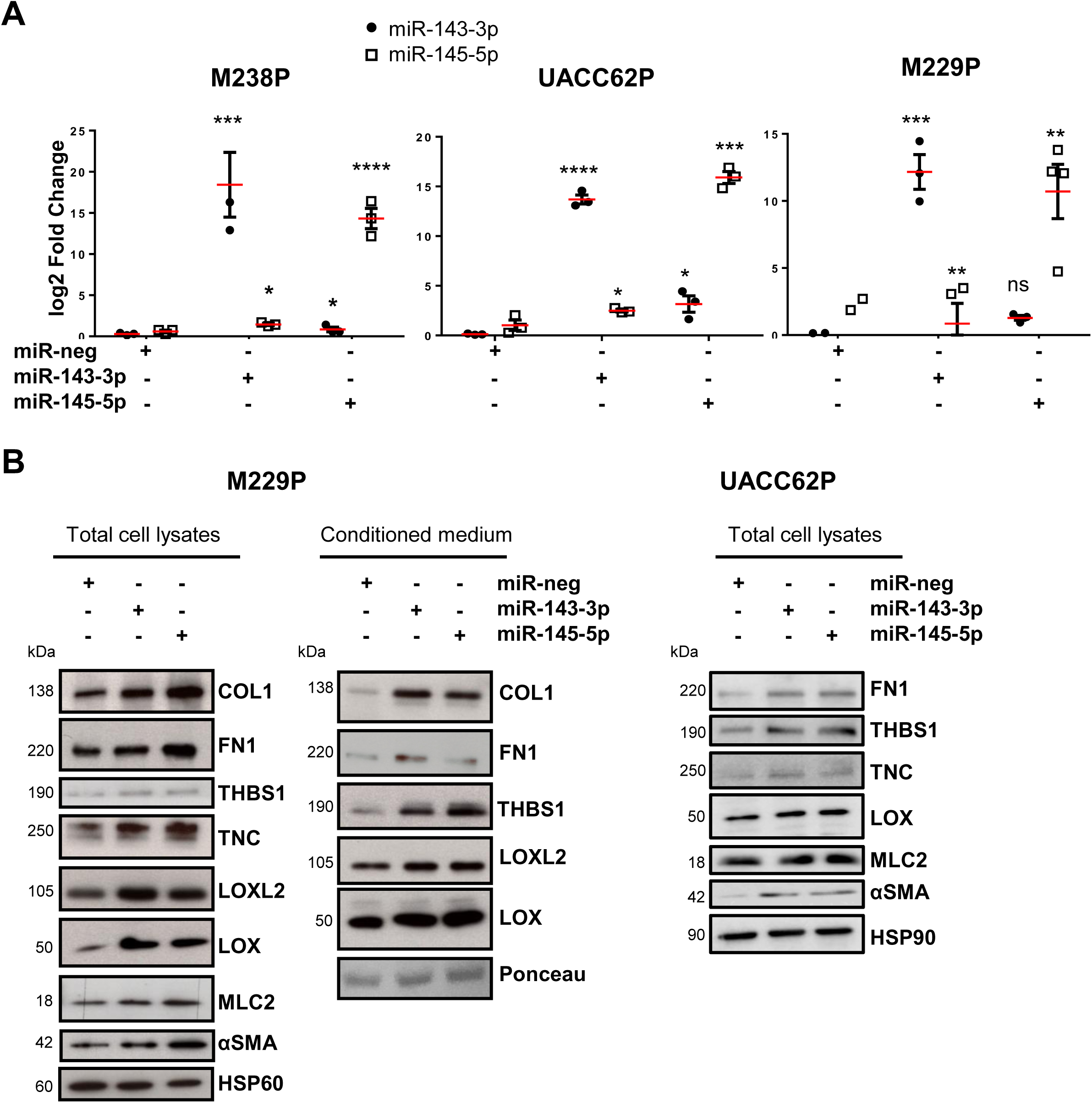
MiR-143/145 cluster plays a role in ECM reprogramming. **(A)** qPCR analysis showing the level of miR-143-3p and miR-145-5p expression after transient transfection of miRNAs mimics (72 h, 30 nM) in 3 distinct melanoma cell lines (M238P, UACC62P, M229P). Data is represented as mean ± SE from a triplicate representative of at least 3 independent experiments. Paired Student t-test has been used for statistical analysis. *P≤0.05 **P≤0.01 ***P≤0.001 ****P≤0.0001 (**B**) Immunoblot analysis of ECM remodeling markers on total cell lysates (M229P and UACC62P) or conditioned medium (M229P) from parental cells transfected with the indicated mimics.

**Fig. S4.**
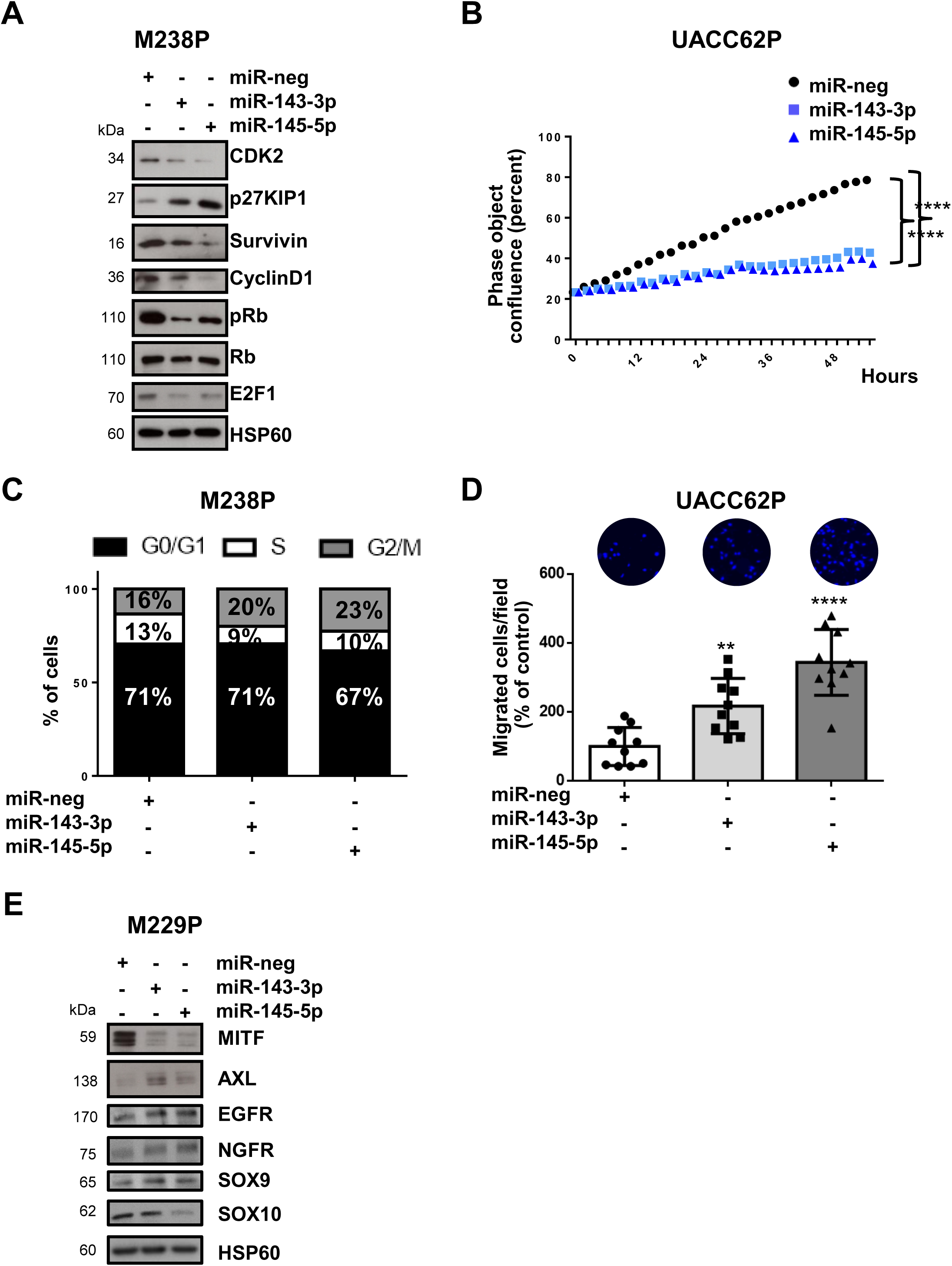
MiR-143/145 cluster drives melanoma cell plasticity and dedifferentiation. Melanoma cells were transfected with control (miR neg), miR-143 or miR-145 mimics (72 h, 30 nM). **(A)** Immunoblot analysis of cell cycle markers on lysates from M238P cells. **(B)** Proliferation curves of parental cells (UACC62P) following transient transfection with miRNA mimics. Time-lapse analysis of cells has been performed with the IncuCyte system. Graph shows quantification of cell confluence. 2-way ANOVA analysis has been used for statistical analysis. ****P≤0.0001. **(C)** Cell cycle distribution of cells cultured in the different conditions. Histograms represent the percentage of cells in different phases of the cell cycle. **(D)** Migration assay of melanoma cells (UACC62P) following transient transfection with miRNA mimics in Boyden chambers. Representative images show migration in control and miR-143-3p or miR-145-5p transfected cells. The histogram represents the quantitative determination of data obtained using ImageJ software. Paired Student t-test has been used for statistical analysis. **P≤0.01, ****P≤0.0001. **(E)** Immunoblot analysis of phenotype switch markers on lysates from parental cells (M229P) transfected with the indicated mimics.

**Fig. S5.**
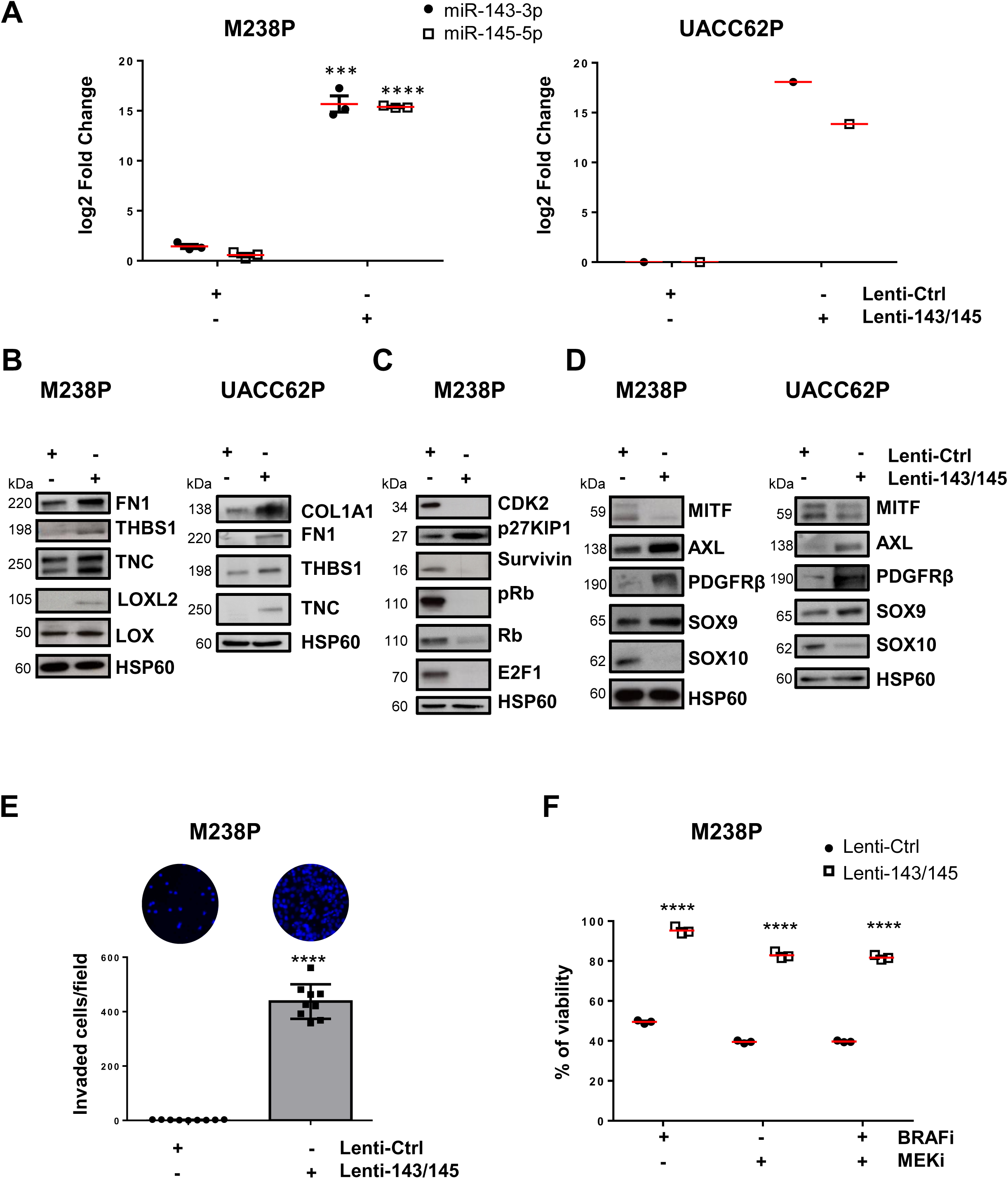
Stable expression of the miR-143/145 cluster promotes ECM remodeling and drives melanoma cell dedifferentiation. **(A)** qPCR analysis showing the level of miR-143-3p and miR-145-5p expression after stable expression following lentivirus transduction of 2 parental cell lines (M238P, UACC62P). Data is represented as mean ± SE from a triplicate representative of at least 3 independent experiments. Paired Student t-test has been used for statistical analysis. *** P≤0.001, ****P≤0.0001. Immunoblot analysis of ECM remodeling **(B),** cell cycle **(C)** and phenotype switch markers **(D)** on total cell lysates from the different stable cell lines. **(E)** Invasion assay in Boyden chambers. Representative images show invasion in control and miR-143/145 expressing cells (M238P). The histogram represents the quantitative determination of data obtained using ImageJ software. Paired Student t-test has been used for statistical analysis. ****P≤0.0001. **(F)** Viability of M238P cells transduced with a control or a miR-143/145 cluster lentivirus was assessed by crystal violet staining upon MAPKi treatment (6 days, BRAFi, Vemurafenib 3 µM, MEKi, Trametinib, 3 µM or BRAFi + MEKi 5 µM). Paired Student t-test has been used for statistical analysis. ****P≤0.0001.

**Fig. S6.**
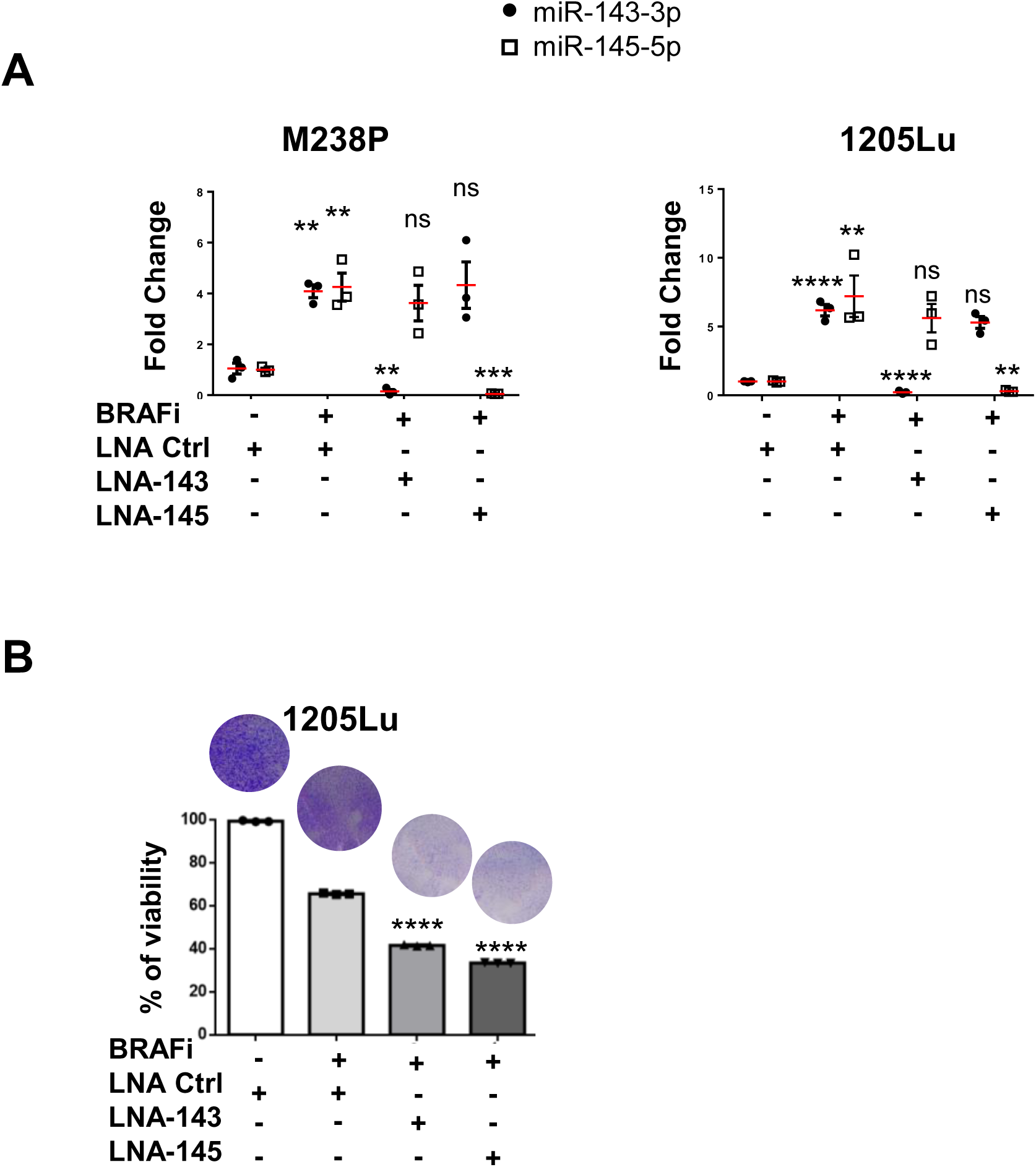
miR-143 and miR-145 inhibition reverses the adaptive response of melanoma cells to MAPK pathway inhibition. Cells were treated with BRAF inhibitor (BRAFi, Vemurafenib) (1 μM, 72h) in the presence or the absence of anti-miR inhibitors (50 nM, 72 h). **(A)** RT-qPCR analysis of miR-143-3p and miR-145-5p expression in parental cells (M238P and 1205Lu) treated with the different combinations of inhibitors. Data is represented as mean ± SE from a triplicate representative of at least 3 independent experiments. Paired Student t-test has been used for statistical analysis. **P≤0.01, ***P≤0.001, ****P≤0.0001. **(B)** Crystal violet viability assay of 1205Lu cells treated with the indicated combinations of inhibitors. Data is represented as mean ± SD from a triplicate representative of at least 3 independent experiments. One-way ANOVA has been used for statistical analysis. ****P≤0.0001.

**Fig. S7.**
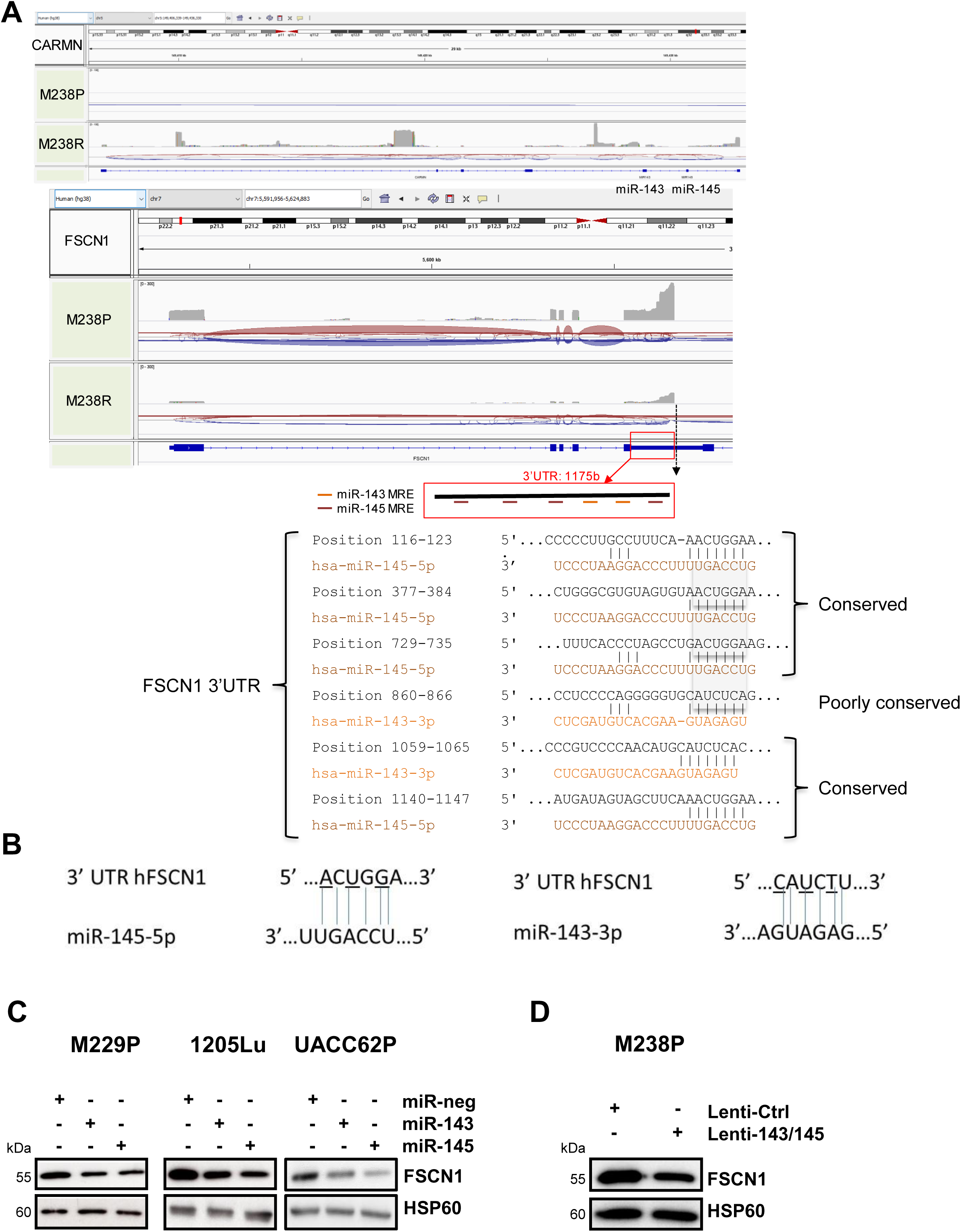
Characterization of the transcripts produced from the miR-143/145 cluster and FSCN1 loci in parental and mesenchymal resistant melanoma cells. **(A)** Screenshot from Integrative Genomic Viewer (IGV) displaying nanopore long-reads RNA-Seq data of the miR-143/-145 cluster / CARMN and FSCN1 loci in M238P and M238R cells (dataset 3, n=1). A strong increase of total reads associated with the CARMN transcripts in M238R compared to M238P cells is shown while the FSCN1 transcript shows the opposite pattern. The red box highlights the FSCN1 3’UTR containing 2 and 4 predicted sites for miR-143-3p and miR-145-5p, respectively. The sequence, pairing and conservation are shown for each predicted site. **(B)** Sequence of h*FSCN1* 3’UTR miR-143 or miR-145 recognition elements and pairing with miR-143 or miR-145 seeds. Bases mutated in the plasmid used for luciferase assay are underlined. **(C-D)** Western Blot analysis of FSCN1 expression in parental cells (M229P, 1205Lu, UACC62P) transfected with the indicated miRNA mimics (72 h, 30 nM) (**C**) and in parental cells (M238P) transduced with the indicated construct **(D)**.

**Fig. S8.**
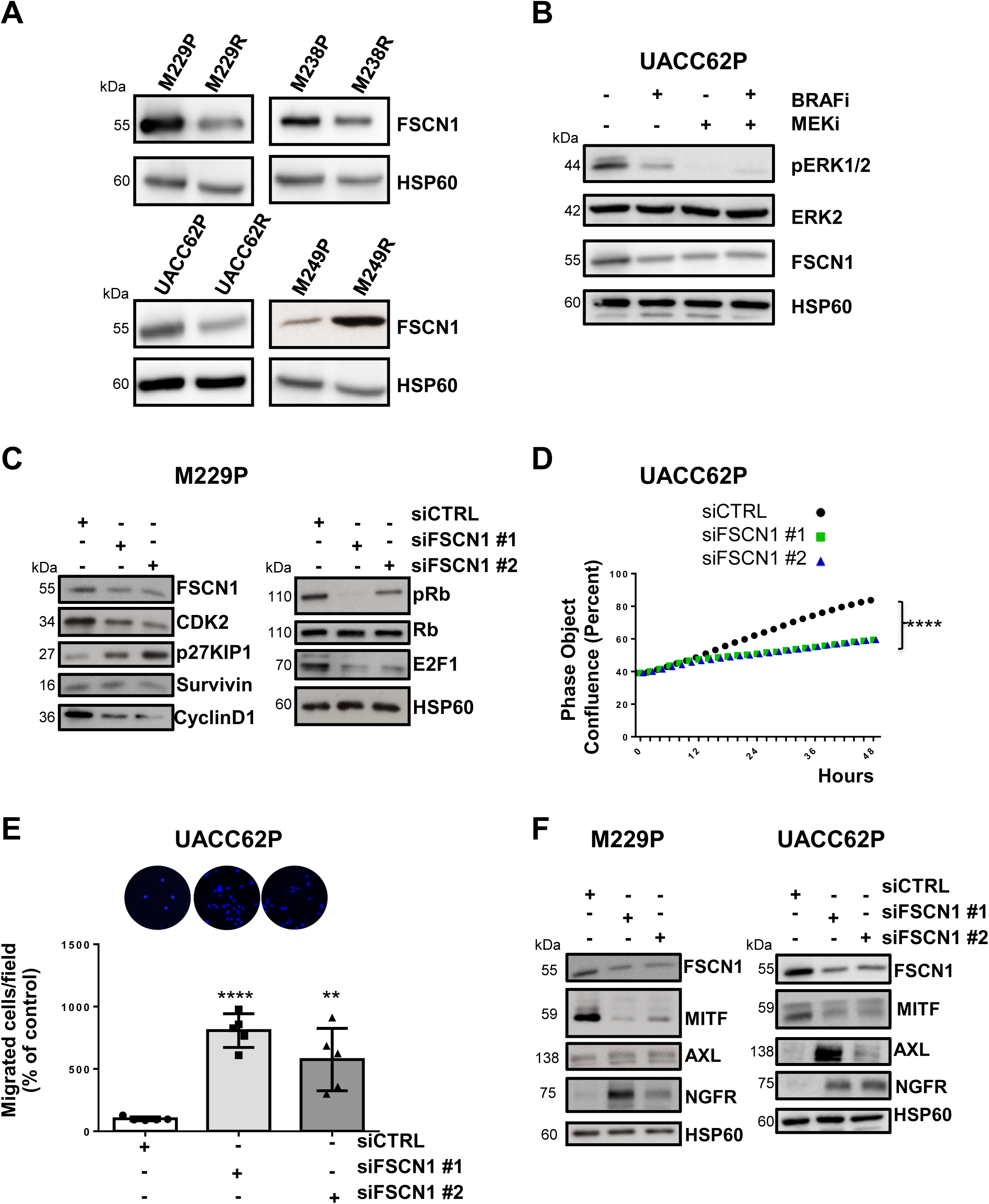
FSCN1 is a functional miR-143/145 target contributing to the phenotypic switch towards an invasive dedifferentiated state. **(A)** Western Blot analysis of FSCN1 expression in parental and paired resistant cells (M238, UACC62, M229, M249). (**B**) Western Blot analysis of FSCN1 levels in parental cells (UACC62P) treated with MAPK-targeted therapies (BRAFi, Vemurafenib, 3 µM), (MEKi, Trametinib, 1 µM), (BRAFi + MEKi 1 µM) for 72 h. **(C-F)** Cells were transfected with two different sequences of siRNAs vs FSCN1 or with a control siRNA (72 h, 100 nM). **(C)** Immunoblot analysis of cell cycle markers on cell lysates from M229P cells cultured for 72 h following transfection with the different siRNAs. **(D)** Proliferation curves using time-lapse analysis of cells with the IncuCyte system. Graph shows quantification of cell confluence. 2-way ANOVA analysis was used for statistical analysis. ****P≤0.0001. **(E)** Migration assay performed in Boyden chambers. Representative images showing migration of UACC62P cells in the indicated conditions. The histogram represents the quantitative determination of data obtained using ImageJ software. Paired Student t-test has been used for statistical analysis. **P≤0.01 ****P≤0.0001. **(F)** Immunoblot analysis of phenotype-switch markers on cell lysates from cells (M229P and UACC62P) transfected with the different siRNAs.

**Fig. S9.**
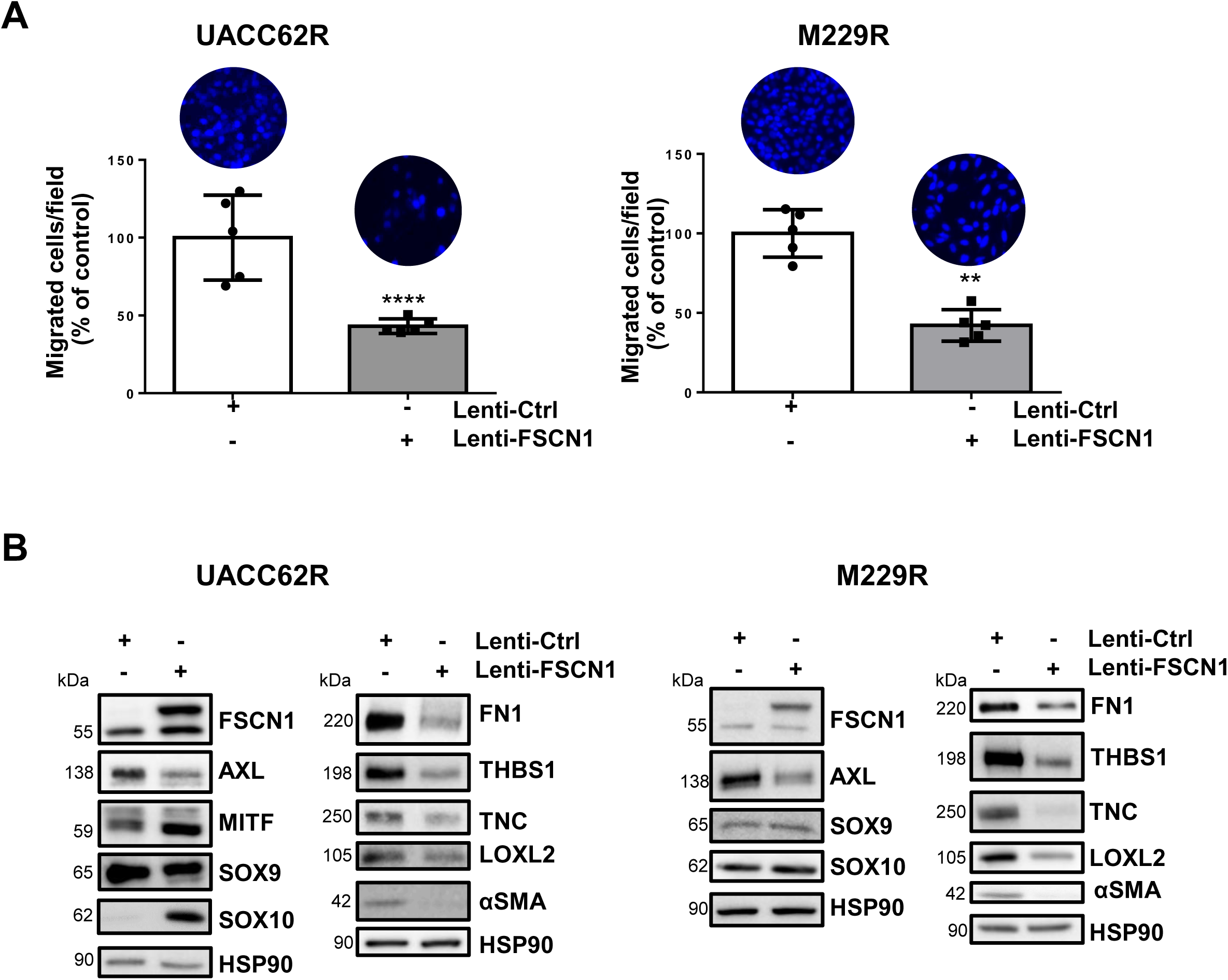
FSCN1 restoration promotes the switch of melanoma cells toward a differentiated cell-state. (A-B) Cells were transduced with a control or a FSCN1 lentiviral construct. **(A)** Effect of FSCN1 overexpression on cell migration (Boyden chambers). Representative images and quantitative determination of data obtained using ImageJ software. Paired Student t-test has been used for statistical analysis. **P≤0.01, ****P≤0.0001. **(B)** Immunoblot analysis of phenotype-switch markers and ECM remodeling markers on cell lysates from control and FSCN1 overexpressing resistant cells (UACC62R, M229R).

**Fig. S10.**
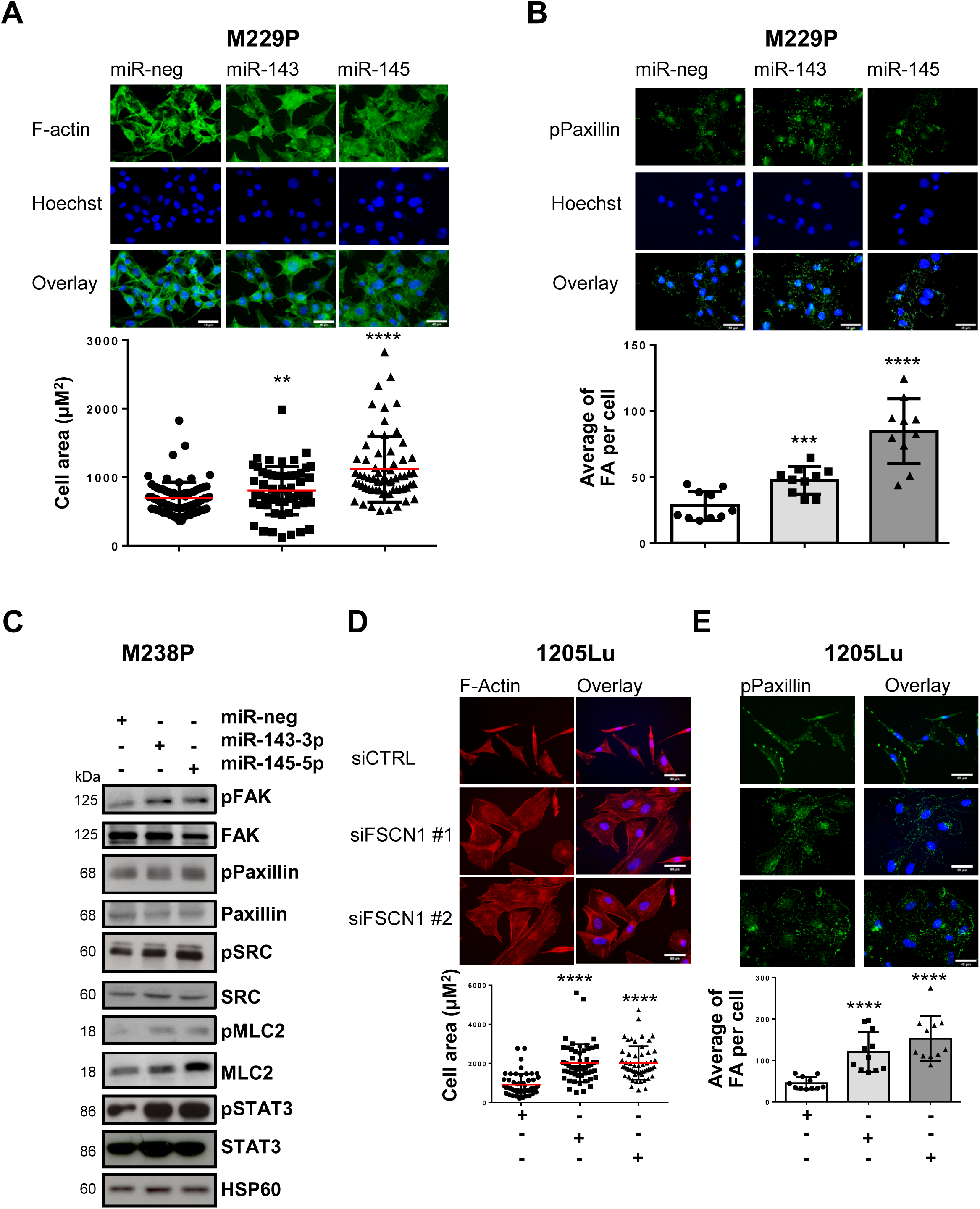
The miR-143-/145 cluster/FSCN1 axis regulates actin cytoskeleton dynamics. (A-C) M229P or M238P cells were transfected with miR-143-3p, miR-145-5p or a control mimic (72 h, 30nM). **(A)** Quantification of cell area in cells stained for F-actin (red) and nuclei (blue). Data is represented as scatter plot with mean ± SD (n≥30 cells per condition). Mann-Whitney U test has been used for statistical analysis. **P≤0.01, ****P≤0.0001. Scale bar 40 μM. **(B)** Quantification of focal adhesions number in cells stained for pPaxillin (green) and nuclei (blue). Focal adhesions number is represented as mean ± SD (n≥30 cells per condition). Each point represents the average number of focal adhesions per cell calculated for each field. Paired Student t-test has been used for statistical analysis. ***P≤0.001, ****P≤0.0001. Scale bar 40 μM. **(C)** Immunoblot analysis of focal adhesion components and cytoskeleton-related pathways in cells transfected with the different mimics. **(D-E)** Cells were transfected with two different sequences of siRNAs vs FSCN1 or with a control siRNA (72 h, 100 nM). **(D)** Quantification of cell area in cells (1205Lu) stained for F-Actin (red) and nuclei (blue). Data is represented as scatter plot with mean ± SD. (n≥30 cells per condition). Mann-Whitney U test has been used for statistical analysis. ****P≤0.0001. Scale bar 40 μM. **(E)** Quantification of focal adhesions number in cells (1205Lu) stained for pPaxillin (green) and nuclei (blue). Focal adhesions number is represented as mean ± SD (n≥30 cells per condition). Each point represents the average number of focal adhesions per cell calculated for each field. Mann-Whitney U test has been used for statistical analysis. ****P≤0.0001. Scale bar 40 μM.

**Fig. S11.**
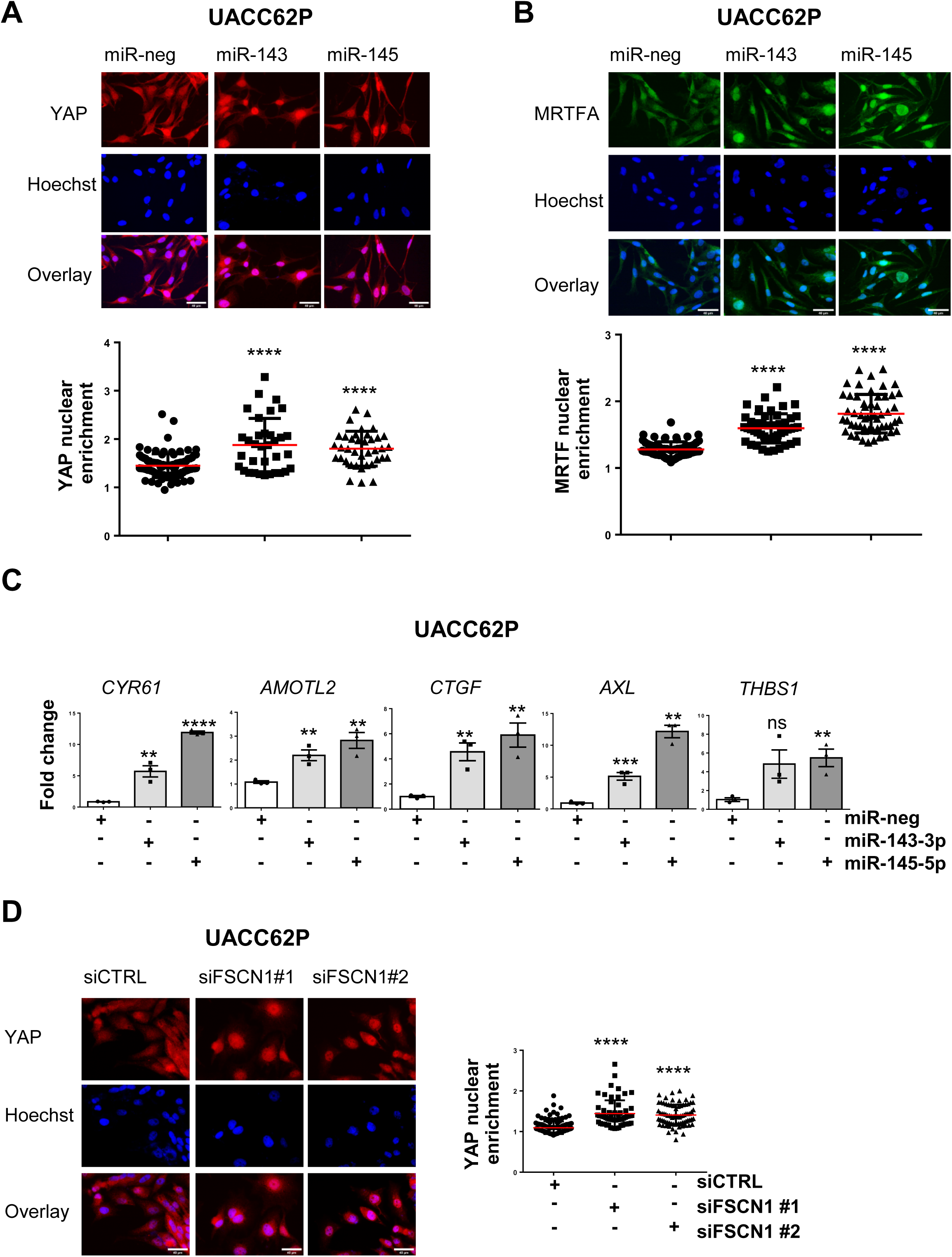
The miR-143-/145 cluster/FSCN1 axis regulates mechanopathways. (A-C) UACC62P cells were transfected with miR-143-3p, miR-145-5p or a control mimic (72 h, 30 nM). **(A)** Effect of miR-143 or miR-145 overexpression on YAP nuclear translocation by immunofluorescence. Cells were stained for YAP (red) and nuclei (blue). **(B)** Effect of miR-143 or miR-145 overexpression on MRTFA nuclear translocation assessed by immunofluorescence. Cells were stained for MRTFA (green) and nuclei (blue). **(A-B)** Data are represented as scatter plot with mean ± SD (n≥30 cells per condition). Mann-Whitney U test has been used for statistical analysis. ****P≤0.0001. Scale bar 40 μM. **(C)** Effect of miR-143 or miR-145 overexpression on the expression of MRTFA/YAP target genes assessed by RT-qPCR. Data are normalized to the expression in control cells. Data is represented as mean ± SE from a triplicate representative of at least 3 independent experiments. Paired Student t-test has been used for statistical analysis.**P≤0.01, ***P≤0.001, ****P≤0.0001. **(D)** Effect of FSCN1 downregulation on YAP nuclear translocation assessed by immunofluorescence in UACC62P cells stained for YAP (red) and nuclei (blue). Data are represented as scatter plot with mean ± SD (n≥30 cells per condition). Mann-Whitney U test has been used for statistical analysis. ****P≤0.0001.

**Fig. S12.**
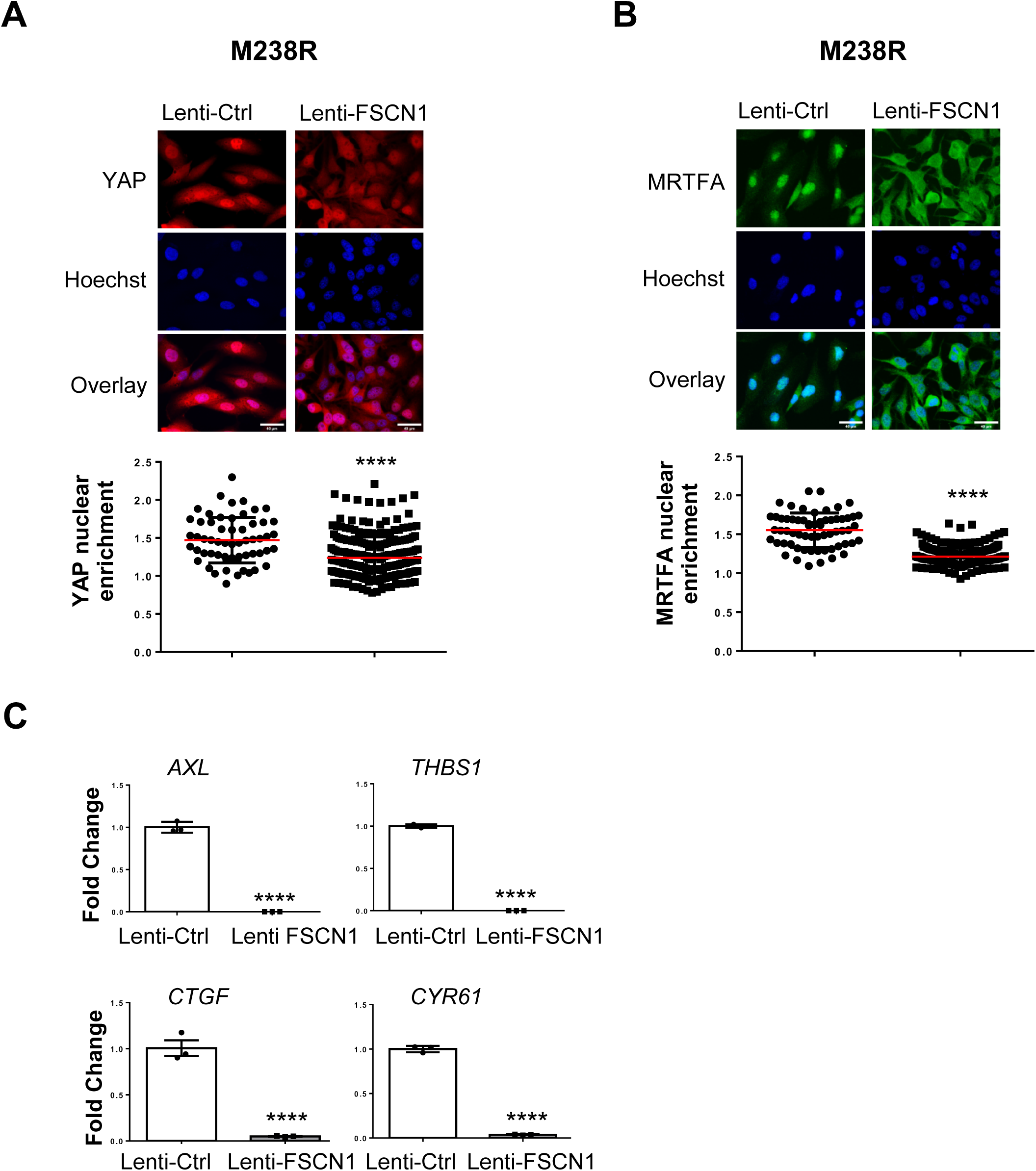
FSCN1 restoration impairs the activation of mechanopathways. BRAFi-resistant M238R cells overexpressing FSCN1 were obtained after transduction with a FSCN1 lentiviral construct. M238R transduced with a Ctrl lentivirus were used as control. **(A-B)** Effect of FSCN1 overexpression on YAP **(A)** and MRTFA **(B)** nuclear translocation assessed by immunofluorescence in cells stained for YAP (red) or MRTFA (green) and nuclei (blue). Data are represented as scatter plot with mean ± SD (n≥30 cells per condition). Mann-Whitney U test has been used for statistical analysis. ****P≤0.0001. Scale bar 40 μM. **(C)** RT-qPCR analysis for the expression of YAP1/MRTFA target genes in M238R cells stably overexpressing FSCN1. Data are normalized to the expression in parental cells. Data is represented as mean ± SE from a triplicate representative of at least 3 independent experiments. Paired Student t-test has been used for statistical analysis. ****P≤0.0001

